# Targeted biomass degradation by the higher termite gut system - integrative omics applied to host and its gut microbiome

**DOI:** 10.1101/2020.02.06.937128

**Authors:** Magdalena Calusinska, Martyna Marynowska, Marie Bertucci, Boris Untereiner, Dominika Klimek, Xavier Goux, David Sillam-Dussès, Piotr Gawron, Rashi Halder, Paul Wilmes, Pau Ferrer, Patrick Gerin, Yves Roisin, Philippe Delfosse

## Abstract

M*iscanthus* sp. is regarded as suitable biomass for different biorefinery value chains. However, due to high recalcitrance, its wide use is yet untapped. Termite is the most efficient lignocellulose degrading insect, and its success results from synergistic cooperation with its gut microbiome. Here, we investigated at holobiont level the dynamic adaptation of a higher termite *Cortaritermes* sp. to imposed *Miscanthus* diet, with a long-term objective of overcoming lignocellulose recalcitrance. We used an integrative omics approach, comprising amplicon sequencing, metagenomics and metatranscriptomics that we combined with enzymatic characterisation of carbohydrate active enzymes from termite gut Fibrobacteres and Spirochaetae. Adaptation to the new diet was evidenced by reduced gut bacterial diversity and modified gene expression profiles, further suggesting a shift towards utilisation of cellulose and arabinoxylan, two main components of *Miscanthus* lignocellulose. Low identity of reconstructed microbial genomes to microbes from closely related termite species, supported the hypothesis of a strong phylogenetic relationship between host and its gut microbiome. Application-wise, this makes each termite gut system an endless source of enzymes that are potentially industrially relevant.

This study provides a framework for better understanding the complex lignocellulose degradation by the higher termite gut system and paves a road towards its future bioprospecting.

## Introduction

In a world of finite biological resources, the agenda of the UN’s Sustainable Development Goals challenges scientific community to develop transformative technologies that would enable the replacement of petroleum-based raw materials and energy with renewable bio-based feedstock. Plant biomass (lignocellulose), being the most abundant and renewable natural resource, could have many applications in different sectors^1^. *Miscanthus* sp. is a rhizomatous grass and owing to its adaptability to various environmental conditions, it shows high potential for sustainable production of lignocellulose over large geographical range^2^. Considering its important agronomic advantages (*e.g.* high biomass yield per hectare, reduced soil erosion and low fertilizer and pesticide requirements), it is suitable for different biorefinery value chains, including bioethanol, biogas, food additives, ingredients for cosmetics, biopharmaceuticals, bioplastics, biomaterials, organic fertilizers and animal feed^3^. Yet, due to the high recalcitrance (resistance of the cell wall components to enzymatic hydrolysis), its use is largely untapped^4^.

In living organisms, enzymatic hydrolysis of lignocellulose is mainly driven by carbohydrate active enzymes (CAZymes^5^). Glycoside hydrolases (GHs) are the primary enzymes that cleave glycosidic linkages. Often, they are assisted by carbohydrate esterases (CEs), polysaccharide lyases (PLs) and other auxiliary enzymes (AAs). With its unique consortium of microorganisms, the termite gut is considered as the most efficient lignocellulose degrading system in nature^6^. Complete loss of gut cellulolytic flagellates in all evolutionary higher termites and acquisition of novel symbiotic bacteria led to improved lignocellulolytic strategies. It allowed for diet diversification from mainly wood-restricted to *e.g.* dry grass and other plant litter, herbivore dung and or soil organic matter at different stages of humification^7^. Until now, most research has focused on endogenous endoglucanases of termites and cellulases from termite gut flagellates^8^. CAZymes from higher termite gut bacteria have only recently started receiving scientific attention^9^.

Here, we investigated the higher termite gut system of *Cortaritermes sp.* (Nasutitermitinae sub-family) from French Guiana savannah. Using 16S rRNA gene amplicon profiling of termite gut bacteria, we demonstrated the adaptation of two termite colonies to *Miscanthus* diet under laboratory conditions. Through the *de novo* metagenomic (MG) and metatranscriptomic (MT) reconstructions, we assessed the distribution of activities within gut community, and we linked it to different bacteria, two main actors being Spirochaetae and Fibrobacteres. Further analysis of gene expression profiles proved microbial functional plasticity (adaptation to changing environmental conditions through differential genes expression), and highlighted the abundance of gene transcripts involved in carbohydrate metabolism and transport. Analysis of the reconstructed community metagenome evidenced CAZyme clusters targeting two main components of *Miscanthus* biomass, namely cellulose and (arabino)xylan. Most of these clusters were allocated to the reconstructed metagenome assembled genomes (MAGs) of Fibrobacteres and Spirochaetae origin. The *de novo* reconstruction of the host epithelial gut transcriptome confirmed its contribution to lignocellulose degradation, and its adaptation to *Miscanthus* diet. Based on the characterisation of purified bacterial CAZymes, we verified the *in silico* predicted activities for many backbone-targeting (*e.g.* endocellulases and xylanases) and debranching enzymes (*e.g.* arabinofuranosidases). To finish, we discussed our findings in the context of enzymes application in the developing biorefinery sector.

## Results

### Structural adaptation of termite gut microbiome to *Miscanthus* diet

To examine enzymatic degradation of *Miscanthus* by the higher termite gut system, two laboratory-maintained colonies (LM1 and LM3) of *Cortaritermes intermedius* were fed exclusively with dried *Miscanthus* straw (Supplementary Fig. 1 and 2). This Nasutitermitinae genus is known to feed on grass tussocks in its natural habitat^10^. Adaptation of the termite gut microbiome (here relative to bacterial communities in termite midgut and hindgut) were monitored during nine months, by high-throughput sequencing the V6-V8 regions of 16 S rRNA gene (Fig. 1a). Quality trimmed reads were assembled into 678 operational taxonomic units (OTUs) assigned to 18 bacterial phyla. Spirochaetae and Fibrobacteres were the most dominant, as previously shown for plant fibre-feeding higher termites (*e.g.* ref.^11^; Supplementary Table 1). By assessing bacterial community structures, we could find clear differences between wild- and *Miscanthus*-adapted microbiomes. Species richness and diversity were also significantly higher (HOMOVA p<0.001) before *Miscanthus* diet was initiated (Fig. 1b). Further application of linear discriminant analysis (LDA) effect size (LEfSe; ref.^12^) demonstrated adaptive selection for specific bacteria (Supplementary Table 2). By analysing food-associated microbes, we estimated the effect of immigration on the termite gut community as negligible (Supplementary Figure 3). All together, these results demonstrated that diet change drives the development of microbial consortium in a unique manner, yet food-associated microbes cannot compete with highly specialised termite gut microbiota for a niche.

**Fig. 1:**
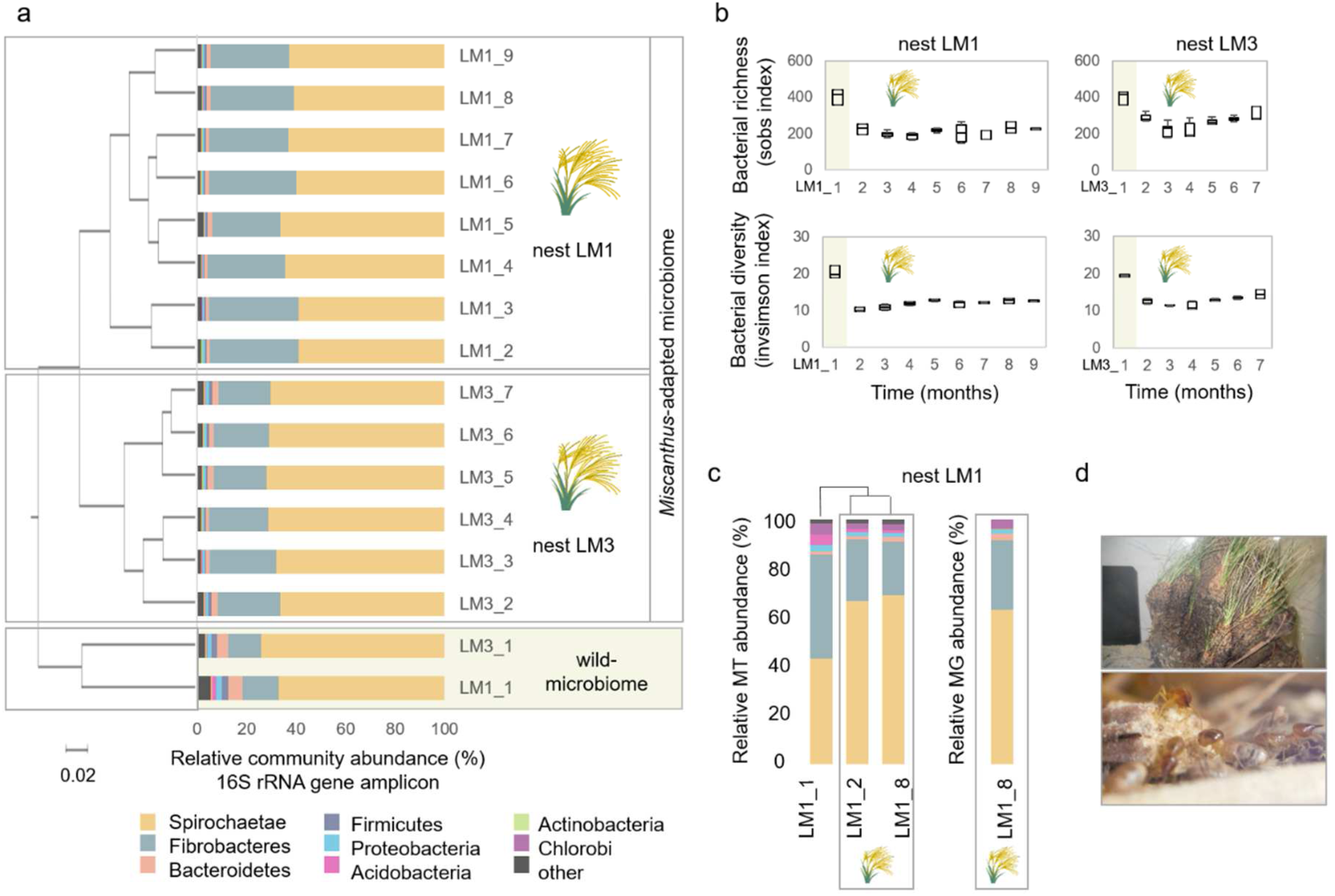
Structural composition of *Cortaritermes* sp. gut microbiome under *Miscanthus* sp. diet. (a) Clustering of samples based on the calculated Bray-Curtis index and phylum level taxonomic assignment of sequencing reads from the 16S rRNA gene amplicon study. (b) Bacterial richness and diversity indices before (highlighted in yellow on sub-figures a and b) and under *Miscanthus*. (c) Relative metatranscriptomic (MT) and metagenomic (MG) reads abundance assigned at the phylum level. Taxonomic gene and gene transcript assignments were inferred from the metagenomic contigs binning and phylum-level bin classification. (d) *Cortaritermes* sp. colony (top) and termite workers under the protection of soldiers while feeding on *Miscanthus* fibres in laboratory conditions (bottom).

### Comparison of *de novo* metatranscriptomic and metagenomic reconstructions

The *de novo* MT was applied to nest LM1 and co-assembling reads from three samples (LM1_1, LM1_2 and LM1_8) yielded 603,579 open reading frames (ORFs), mainly representing partially reconstructed gene transcripts. The *de novo* MG reconstruction was also applied to colony LM1 at sampling point LM1_8 and it yielded 211,724 ORFs annotated on 64,347 contigs for the total assembly size of 177,5 million base pairs (Mbp). For both datasets, initial public database-dependent taxonomic classification of genes and gene transcripts pointed to the abundance of Firmicutes (Supplementary Fig. 4), what contrasted the results of community structure analysis. Subsequent binning of MG contigs and phylum-level annotation of the resulting bacterial bins allowed assigning correct taxonomic origin, confirming the metagenomic abundance of Spirochaetae and Fibrobacteres (Fig. 1c; Supplementary Fig. 5). Following mapping of RNA reads to the MG assembly (referred to as “RNA-seq” analysis), we could confirm transcriptional dominance of these two bacterial phyla as well. Incomplete public databases and extensive horizontal gene transfer were previously proposed as the origin of this misclassification^9^.

Based on the classification of genes and transcripts to broad functional categories such as KEGG ontology profiles (KOs), congruency between the *de novo* MG and MT reconstructions was high (Supplementary Fig. 6). However, out of the *de novo* MT reconstructed gene transcripts of prokaryotic origin only 37.8 % showed significant similarity to the *de novo* MG genes (blast e-value≤10^−5^), sharing on average 76.04 % of amino acid identity (Supplementary Fig. 4). Coherent to our study, differences in functional gene profiles between MG and MT reconstructions have been previously underlined^13^. Even in the context of the termite gut, some authors highlighted the value of the *de novo* MT assembly in retrieving highly expressed genes^9,14^.

### Genomic potential and functional adaptation of termite gut microbes to *Miscanthus* diet

Aggregation of 68.9 ±1.8 % of the *de novo* reconstructed gene transcripts into clusters of orthologous genes (COGs) pointed at functional microbiome stability at the different stages of feeding campaign (Supplementary Table 3). Consistently with previous reports^11,14,15^, cell motility and chemotaxis together with carbohydrate transport and metabolism were the two most highly expressed gene categories. Reconstruction of (nearly) complete metabolic modules was quite similar between Fibrobacteres and Spirochaetae (Supplementary Table 5). Further in depth analysis of gene expression profiles evidenced adaption to *Miscanthus* diet (Fig. 2; Supplementary Tables 4 and 5), at the same time identifying several biologically informative features differentiating these two bacterial populations. Both bacterial populations were capable of nitrogen fixation and glycogen synthesis, but the two pathways were enriched in Fibrobacteres. Expression of Amt ammonium transporters was highly up-regulated, and together with increased abundance of gene transcripts involved in urea transport and metabolism (restricted to Spirochaetae), it indicated nitrogen deficiency of a *Miscanthus* fed termite colony. Both Spirochaetae and Fibrobacteres could also synthetize ten essential amino acids that animals cannot synthetize *de novo.* Even though, nitrogen provisioning by bacterial symbionts is not employed by all herbivorous insects, this strategy was proposed as a mechanism contributing to the success of termites^14^ and herbivorous ants^16^ in their marginal dietary niches. Importance of lignocellulose degradation under *Miscanthus* diet was evidenced by increased abundance of transcripts broadly assigned to cellulose and xylan processing KOs (Fig. 2; Supplementary Table 4). Multiple sugar ABC transporters were up-regulated in the Spirochaetae metatranscriptome, while they were nearly absent from the MG and MT reconstructions of Fibrobacteres origin. This observation could point to the governance of exogenous carbohydrates uptake and utilisation by Spirochaetae.

**Fig.2:**
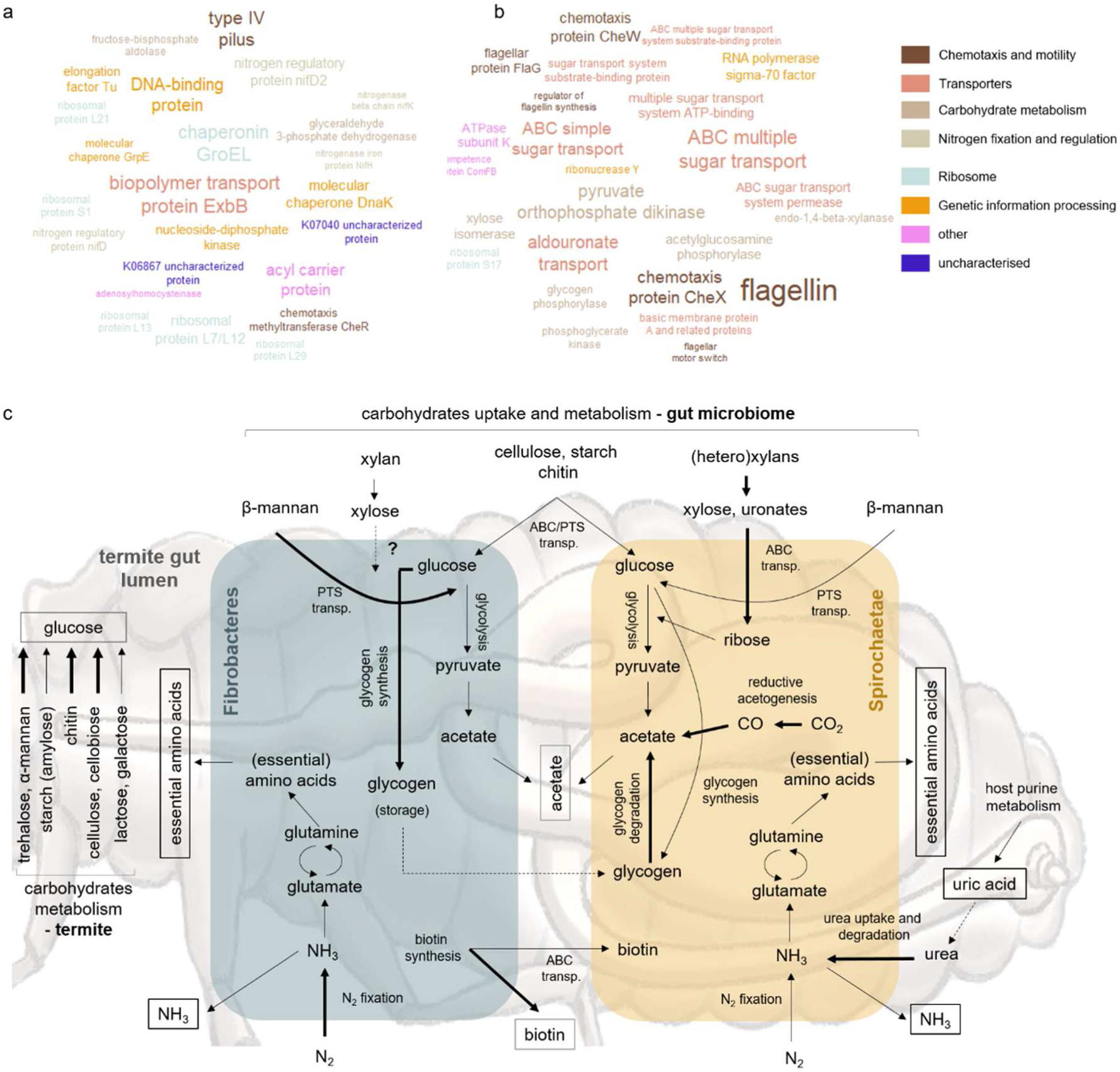
Functional characterisation of the termite gut system feeding on *Miscanthus* sp. (a, b) Tag clouds of enriched (LefSe LDA>2, p<0.05) KOs reconstructed from the *de novo* metatranscriptomics for the termite gut Fibrobacteres (a) and Spirochaetae (b) populations at LM1_8. Top 25 most abundant KOs are displayed. (c) Simplified metabolic reconstruction, with a focus on carbohydrate metabolism, for the termite gut lignocellulolytic system. Hypothetical pathways are indicated with dashed lines. Metabolic pathways enriched in Fibrobacteres and Spirochaetae populations (metatranscriptomes) are indicated with bold lines. Metabolites putatively shared between gut bacteria and the host are indicated with square boxes.

### Diversity and abundance of termite gut microbial CAZymes

The *de novo* MT reconstruction yielded over 2000 of manually curated transcripts assigned as CAZy-coding genes (*cazymes*). Out of these, 38.4 % were further assigned to 55 GH families. The *de novo* MG reconstruction resulted in close to 7000 different *cazymes*, 43.6 % of which were assigned to 86 GH families (Supplementary Fig. 6 and 7). Although there was a good correlation between the distributions of identified glycoside hydrolases to different GH families (Pearson R^2^ 0.83), roughly 150 *cazymes* identified in the *de novo* MT were also reconstructed from the assembled metagenome. Novelty of reconstructed *cazymes* was evidenced through sequence comparison to NCBI refseq database, and a metagenomic dataset previously generated for *Nasutitermes* sp.^15^ (Supplementary Fig. 4). In the latter case, average amino acids identity for the 943 query hits equalled 65.4 ± 19.9 %, pointing to the diversity of microbial *cazymes* in guts of different termite species, even those phylogenetically closely related.

Differential expression of specific *cazymes* at different stages of the feeding experiment evidenced quick and specific adaptation of gut microbes to digest *Miscanthus* (Fig. 3). There were roughly 29.7 % of common *cazymes* transcripts between LM1_8 and control LM1_1 sample, while over 55 % were shared between LM1_8 and LM1_2 (both adapted to *Miscanthus*). Along the experiment, GH5 (mainly subfamilies GH5_4 and GH5_2) was the most highly expressed family. Still, its cumulative expression nearly doubled under *Miscanthus* diet (Fig. 3b). Other abundant families included GH43, GH10 and GH11, all potentially involved in (hetero)xylan degradation. The latter was previously shown as largely expressed by the termite gut fibre-associated Spirochaetae^9^. Following manual curation, we removed three highly abundant but only partially reconstructed GH11 gene transcripts, what reduced initial over-dominance of this CAZy family by 3.3 ±0.9 fold (Supplementary Fig. 8). Similarly, only fragmented GH11 genes were recovered from the reconstructed metagenome. We hypothesise that closely related Spirochaetae strains contain highly similar GH11 genes, possibly shared by horizontal gene transfer, likewise it was shown for *e.g.* human gut Bacteroidetes^17^. Their conserved nature may impede proper gene reconstruction from sequencing reads, similarly to other structurally conserved genes such as 16S rRNA or transposase, that typically do not reconstruct into larger genomic fragments^18^. GH45 was only present in Fibrobacteres and it was the second most expressed GH family under *Miscanthus* diet. According to the CAZY database, nearly 95 % of proteins assigned to this family are of eukaryotic origin (Supplementary Fig. 10), and show endocellulase activity. Previously, GH45 CAZymes were characterised from lower termite symbiotic protists^19^. Lytic polysaccharide monooxygenases (LPMOs) typically assigned to AA9 (fungal) and AA10 (predominantly bacterial enzymes) families^20^ were neither present in the reconstructed metagenome nor metatranscriptome.

**Fig. 3:**
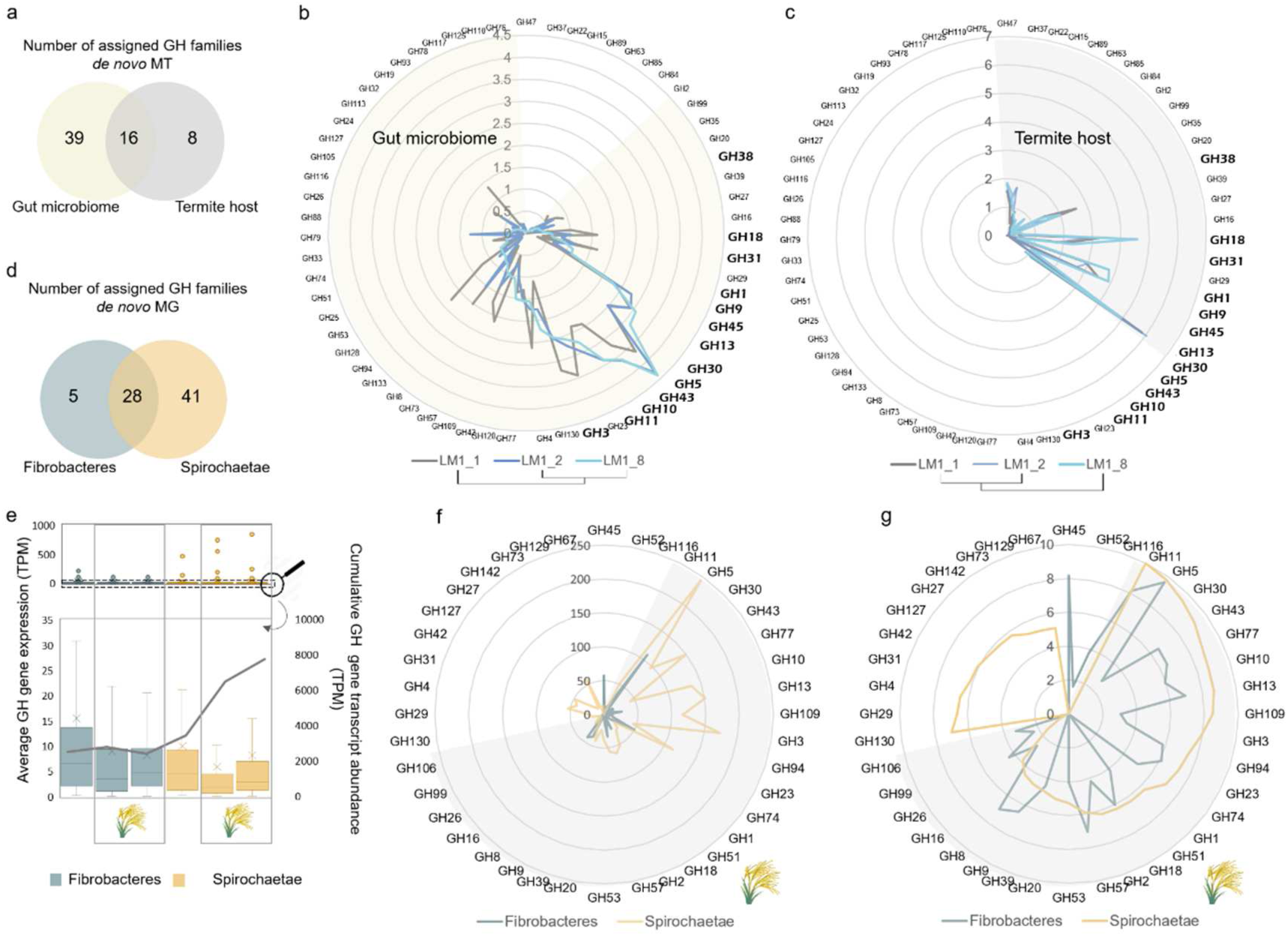
Carbohydrate active enzymes (CAZymes) reconstructed from metatranscriptomic (MT) and metagenomic (MG) reads for the termite gut system. (a) Venn diagram showing the number of assigned glycoside hydrolase (GH) families for the termite gut microbiome and host gut epithelium. (b,c) Comparison of the gene expression profiles (*de novo* MT, log2 transformed) for gene transcripts assigned to the different GH families and at the different stages of the *Miscanthus* feeding experiment, for the gut microbiome (b) and the host gut epithelium (c). (d) Venn diagram showing the number of assigned GH families for Fibrobacteres and Spirochaetae, based on the *de novo* MG reconstruction. (e) Average CAZyme genes expression and cumulative gene expression at the different stages of the feeding experiment and analysed separately for the Fibrobacteres and Spirochaetae populations. Lower panel is a zoom on the gene expression profiles with the outliers (highly expressed genes; in some cases representing only partially reconstructed genes) removed. (f, g) Number of genes (f) and cumulative abundance of the most abundant GH families (g, RNA-seq log2 transformed) at the time point LM1_8 (the end of the *Miscanthus* feeding experiment), and visualised separately for the Fibrobacteres and Spirochaetae populations. Transcripts abundance (g) is calculated based on the RNA mappings (RNA-seq) to the MG contigs. Shaded parts correspond to shared GH families.

### Expression and activities of GHs from Fibrobacteres and Spirochaetae

Based on the RNA-seq analysis, Fibrobacteres and Spirochaetae expressed respectively 47.9 ±14.8 % and 45.6 ±18.5 % of their *cazymes* genomic content when the termite was fed with *Miscanthus*. Total diversity of Spirochaetae *cazymes* was 2.3 ±0.5 fold higher than for Fibrobacteres (Fig. 3d-g). Their cumulative transcriptional abundance was also higher for Spirochaetae, however calculated average gene expression was slightly higher for Fibrobacteres. This observation was consistent across different GH families (Supplementary Fig. 11). As various GH families are characterised with broad functionalities, using peptide-based functional annotation^21^ we further assigned *in silico* specific functions (EC numbers) to 60.3 ±2.2 % of gene transcripts classified as GHs. In many cases, these predictions were experimentally validated (Supplementary Table 6). We confirmed β-glucosidase, endocellulase, endoxylanase and arabinofuranosidase activities of several Spirochaetae CAZymes. We also characterised active endoxylanases and endomannanases from Fibrobacteres.

Abundance of transcripts associated with endocellulase (EC:3.2.1.4) and endoxylanase (EC:3.2.1.8) increased under *Miscanthus* diet (for both Fibrobacteres and Spirochaetae), while those involved in chitin and starch (α-glucans) degradation decreased (Fig. 4, Supplementary Fig. 12). Endocellulase-assigned transcripts were nearly equally abundant between Fibrobacteres and Spirochaetae, while abundance and diversity of endoxylanases of Spirochaetae origin was much higher. Most of the assigned endocellulases were classified as GH5_4 enzymes (Supplementary Fig. 8.). Phylogenetic reconstruction comprising the previously characterised CAZymes from this family revealed the presence of multiple protein clusters separately grouping Spirochaetae and Fibrobacteres GHs (Fig. 5a). Concurrent inspection of reconstructed genomic fragments suggested the existence of different *cazymes* loci containing GH5_4 genes (Supplementary Fig. 13). Interestingly, CAZymes previously characterised to possess single enzymatic activity (mostly endocellulase and to a lower extent endoxylanase) grouped in upper part of the tree. Lower part of the tree mainly contained multi-functional enzymes (single enzyme simultaneously acting on cellulose and xylan). Suggested enzymatic multi-functionality was further confirmed for a selected GH5_4 CAZyme representing Spirochaetae cluster IX. Purified protein was shown to be an endocellulase acting on carboxymethylcellulose (CMC) and glucomannan (Fig. 5b). In addition, activity on xylan and arabinoxylan was confirmed. This gene was also one of the most highly expressed *cazymes* under *Miscanthus* diet, hypothesising the importance and interest for bacteria to express multi-functional enzymes. To our best knowledge, it represents first GH5_4 CAZyme of Spirochaetae origin ever characterised, and first multi-functional enzyme of higher termite gut prokaryotic origin.

**Fig. 4:**
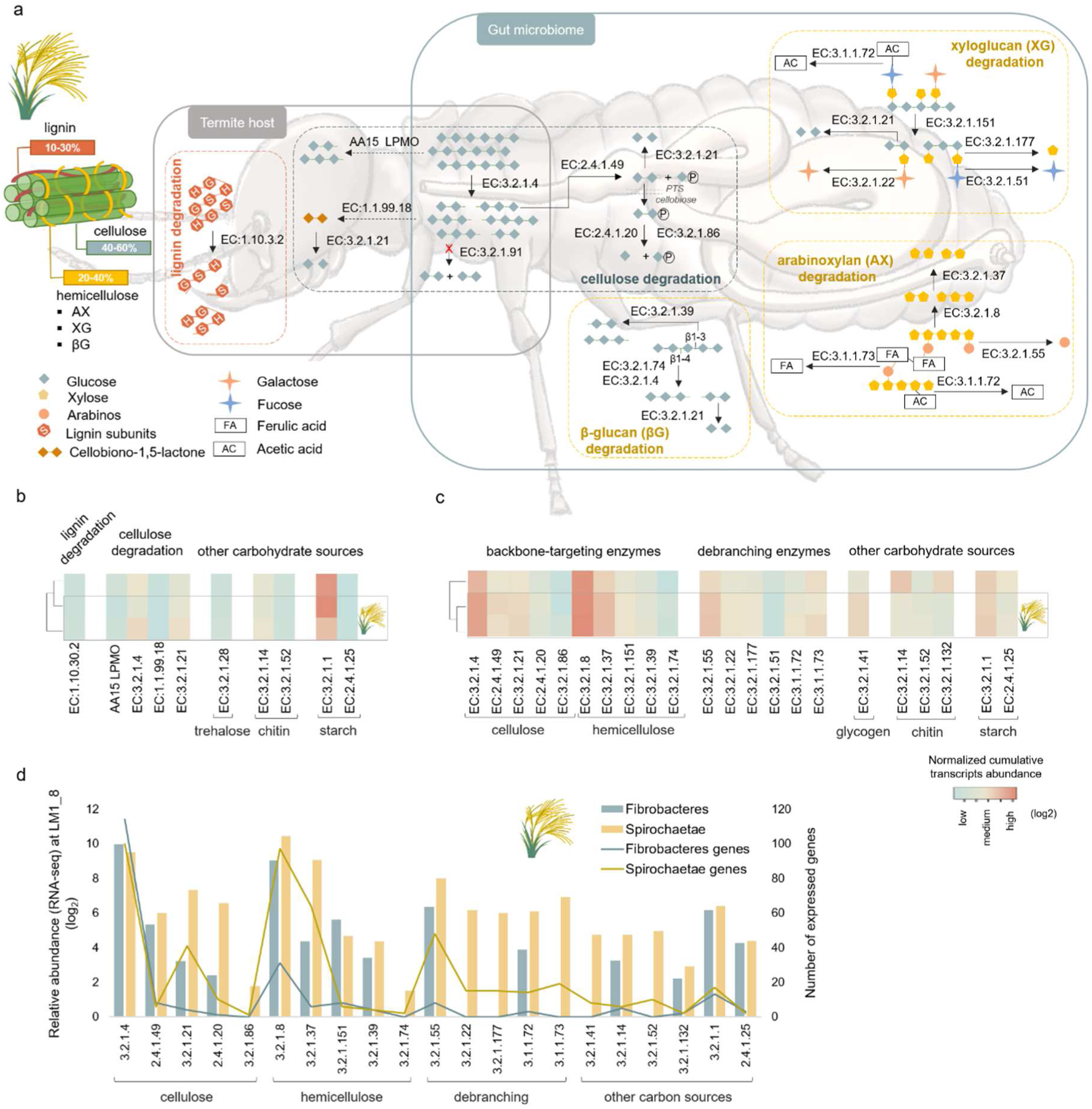
Characterisation of the termite gut lignocellulose degradation strategies. (a) Simplified overview of enzymatic pathways involved in the degradation of main components of the *Miscanthus* biomass, based on enzymes (gene transcripts assigned an EC number) revealed in our study. Dashed lines indicate hypothetical pathways. CAZymes gene expression profiles (*de novo* MT) at the different stages of the *Miscanthus* feeding experiment analysed for the termite gut epithelium (b) and the termite gut microbiome (c). Gene expression analyses were done separately for the termite gut epithelium and the gut microbiome, therefore data on sub-figures b and c should only be compared within a single sub-figure. (d) Relative CAZymes gene transcripts abundance (RNA-seq) and gene numbers assigned to different enzymatic categories and analysed separately for Fibrobacteres and Spirochaetae populations for LM1_8 sample.

**Fig. 5:**
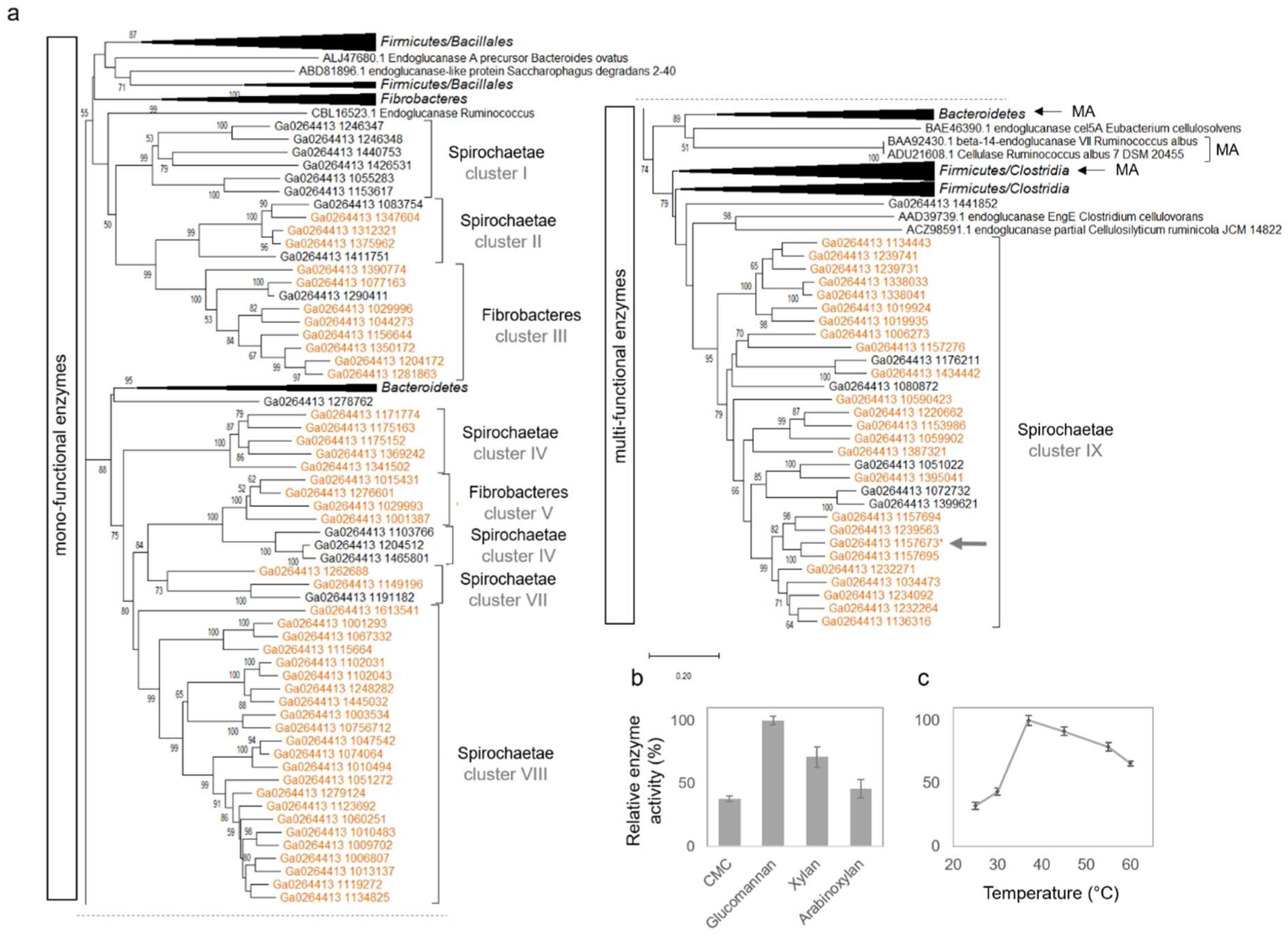
Characterisation of the GH5_4 family. (a) Unrooted neighbor-joining tree containing the *de novo* reconstructed genes from MG study (genes expressed under *Miscanthus* diet are highlighted in orange on the tree). Tree was cut in two parts along the dashed line. All GH5_4 characterised proteins were retrieved from the CAZY database and included on the tree. Clusters indicated with an arrow and designated as “MA” contain known multi-functional enzymes. The percentage of replicate trees in which the associated sequences clustered together in the bootstrap test (500 replicates) are shown next to the branches. Final alignment involved 157 amino acid sequences. Protein from the Spirochaete*s* cluster IX indicated with a grey arrow was heterologously produced and characterised. (b) Activity profiles for the heterologously produced and purified protein tested against CMC, glucomannan (galactomannan was negative, data not shown), xylan and arabinoxylan. (c) Optimal temperature was assessed for glucomannan substrate.

### MAGs reconstruction and carbohydrate utilisation gene clusters

Reconstruction of draft microbial genomes enriched in *Miscanthus*-adapted termite gut microbiome resulted in 20 MAGs with completeness above 50 % and contamination below 10 %, including eight Fibrobacteres-assigned MAGs, six Spirochaetae (four of Treponema origin) and three novel MAGs representing Proteobacteria phylum (Supplementary Table 7 and Supplementary Fig. 14). Average nucleotide identity (ANI) to MAGs from gut microbiomes of several higher termite species^22^ equalled roughly 77.1 ±1.8 %, confirming the novelty of our MAGs. Frequency of *cazymes* was higher in the Treponema genomes and they were also enriched in GHs. Over 36 % of annotated *cazymes* were arranged into 1096 gene clusters containing more than one CAZy encoding gene. Similar gene clusters were recently discovered in gut microbiota of a wood-feeding termite *Globitermes brachycerastes*^23^. Putative cellulose-utilisation gene clusters were the most highly expressed in Fibrobacteres MAGs, suggesting its major contribution to cellulose degradation (Fig. 3 and 4). Endo-xylanase-encoding genes were often co-localised with endocellulases, however xylosidases were scarce, questioning the ability of Fibrobacteres to utilise complex xylans. Reconstructed Spirochaetae genomes were enriched in cazymes clusters targeting among others alpha glucans, (arabino)xylans (AX), beta glucans, cellulose, chitin, galactans, mannans and xyloglucans. Largely complete AX-targeting clusters were assigned to MAG_17 and MAG_1, both representing Treponema. All of them were transcriptionally up-regulated under *Miscanthus* diet (Supplementary Fig. 15), what is consistent with high AX content of *Miscanthus* hemicellulose. Genes encoding for carbohydrate transporters (Supplementary Table 8) and in some cases even the two-component system response regulators and chemotaxis were adjacent to several *cazymes* clusters.

Organisationally similar to polysaccharide utilization loci (PULs) systems employed by Bacteroidetes^24^, *cazymes* clusters reconstructed in our study more resemble the concept of Gram-positive PULs recently proposed by Sheridan et al., (2016) in the context of human gut Firmicutes^25^. Taken into account high sequence similarity of Spirochaeteae and Firmicutes *cazymes*, it is possible that whole *cazymes* clusters were acquired from Firmicutes in the course of evolution. Although not yet shown for Spirochaetae, Bacteroidetes PULs are often encoded within integrative and conjugative elements enabling their transfer among closely related species^17^.

### Host functional adaptation to *Miscanthus* diet

Following taxonomic annotation, 16,416 gene transcripts of eukaryotic origin were classified to *Arthropoda.* Based on the presence of 271 conserved orthologous reference eukaryotic genes^26^, we estimated the completeness of our termite gut epithelial transcriptome at 89.5 %. Number of assembled ORFs was in line with the two published termite genomes, *Macrotermes natalensis* (16,140 protein-coding genes; ref.^27^) and *Zootermopsis nevadensis* (15,459 protein coding-genes; ref.^28^; Supplementary Table 9 and Supplementary Fig. 16). Similarly to gut microbiome, gene transcripts related to carbohydrate transport and metabolism were abundant in the host transcriptome, showing importance of carbohydrates metabolism to the termite lifestyle (Supplementary Table 3). Over 9,000 transcripts were assigned a KO number and a summary of complete and partially reconstructed metabolic pathways is provided in Supplementary Table 4. To our knowledge, this reconstruction represents first transcriptome of a higher termite form Nasutitermitinae family.

We further identified 270 transcripts containing CAZY modules assigned to four main classes (GH, CE, AA and GT) and associated CBMs. Glycoside hydrolases were encoded by 97 genes, and their diversity patterns were similar to those identified in *M. natalensis* and *Z. nevadensis* (Supplementary Table 9). They were assigned to 24 different GH families, out of which eight were not represented in the gut microbiome (Fig. 3a, c). The highest transcriptional abundance was attributed to GH13 (typically assigned as alpha glucanases; Supplementary Fig. 17), and it slightly decreased only towards the end of *Miscanthus* campaign. Based on rather constant expression profiles of chitodextrinase, chitin utilization by the host did not change significantly upon *Miscanthus* feeding (Supplementary Fig. 9). This indicates constant complementation of diet with nitrogen rich chitin, originating from either necrophagy and/or cannibalism, or fungi-colonised food stored in the nest, as previously proposed for other higher termite species^29^.

Transcriptional abundance of endocellulases increased at later stages of *Miscanthus* feeding, suggesting a shift towards increased cellulose utilization by host. Interestingly, one gene assigned to AA3 family was classified as putative cellobiose oxidoreductase (EC 1.1.99.18; Fig. 4a, b). This type of oxidase is involved in oxidative cellulose and lignin degradation in wood-decaying fungi^30^. Moreover, two other transcripts assigned to GH45 family (putative endoglucanases) were present in the reconstructed transcriptome, and their expression slightly increased under *Miscanthus* diet. More stringent homology search revealed their closest similarity to *Anaeromyces contortus*, sp. nov. (Neocallimastigomycota), an anaerobic gut fungal species recently isolated from cow and goat faeces^31^. In insects, GH45 *cazymes* were previously detected in the genomes of *Phytophaga* beetles^32^ and until now they were not reported from sequenced termite genomes. In addition, one gene transcript sharing 54 % of identity (at protein level) with a newly characterised AA15 LPMOs from *Thermobia domestica*^33^ was also identified. This insect has a remarkable ability to digest crystalline cellulose without microbial assistance. Further blast analysis revealed a homologous gene in the genome of *Z. nevadensis*, suggesting that next to certain eukaryotes (*e.g.* crustaceans, molluscs, chelicerates, algae, and oomycetes) termites might be able to oxidatively cleave glycosidic bonds.

### Diet on *Miscanthus*: who does what?

*Miscanthus* biomass is mainly composed of cellulose (41.4 ±2.9 %), hemicelluloses (25.8 ±5.2 %) including arabinoxylans, xyloglucans, β-glucans, and lignin (21.4 ±3.6 %) and other trace components^34^. Based on the annotation of GH profiles, termite on its own can digest amylose (starch), cellulose and/or cellobiose, lactose, galactose, chitin, mannan and mannose (mainly α-mannan presumably contained in fungus cell wall and to a lower extent hemicellulose β-mannans), trehalose, mammalian glycans (*e.g.* N-acylsphingosine) and bacterial cell wall components (Fig. 4a, b). Hemicellulose degradation is exclusively conducted by gut microbes, and no putative hemicellulolytic genes were recovered from the reconstructed termite transcriptome. Slightly increased transcriptional abundance of laccase-coding gene might indicate termite ability to target lignin. Beyond relatively high lignin content, recalcitrance of *Miscanthus* biomass is mainly enhanced by other features, in particular acetylation and esterification with ferulic acid (FA). While acetyl esterase activity can be deduced from Fibrobacteres and Spirochaetae metatranscriptomes, the ability to break ferulic linkages seems limited to Spirochaetae. Putative feruloyl esterase from CE1 family are contained within AX-targeting *cazymes* clusters in Spirochaetae MAGs, and were all up-regulated under *Miscanthus* diet (Supplementary Fig. 15).

Based on the diversity and expression patterns of GHs, different sugar transporters (mainly ABC and to a lesser extent PTS; Supplementary Table 8) and specific sugar isomerases, Spirochaetae population is able to utilise a wider range of sugars (including glucose, glucoronate, rhamnose, arabinose, mannose, xylose, ribose and fucose) than Fibrobactere*s* (mainly glucose, mannose and possibly ribose). Both bacterial populations can target the backbone of cellulose, xylans and mannans (the latter is not abundant in *Miscanthus* biomass). Enrichment of Fibrobacteres metatranscriptome in endoglucanases (both targeting 1–3 β and 1–4 β glycosidic bonds as present in β-glucans and cellulose, respectively) shows its preference for carbohydrates with a glucose-unit backbone. Fibrobacteres also express endoxylanases, and we could confirm experimentally xylanase activity for one GH11 enzyme (Supplementary Table 6). However, hardly represented xylosidase-assigned gene transcripts and the absence of any xylose transporters and other known genes involved in xylose utilisation, would question the ability of Fibrobacteres to utilise xylans. Co-localisation of many endoxylanases together with potential endocellulases in the reconstructed Fibrobacteres MAGs further suggest that termite gut Fibrobacteres mainly remove xylan polymers from *Miscanthus* fibres to better expose cellulose to the action of own endocellulases (Supplementary Fig. 13). By contrast, xylose isomerases and xylulose kinases were enriched in Spirochaetae metatranscriptome and both were highly expressed under *Miscanthus* diet. All reconstructed gene transcripts were assigned to Spirochaetae, and together with enrichment of endoxylanase transcripts, it confirms the ability of these bacteria to degrade xylans.

Based on the *in silico* prediction of enzyme sub-cellular localizations, most of the endoxylanases from Fibrobacteres and Spirochaetae are exported outside the cell (Supplementary Table 10), suggesting initial degradation of xylan backbone in the extracellular space. Many Spirochaetae endocellulases are also putative extracellular enzymes or anchored to the outer membrane. In contrast, multiple Fibrobacteres endocellulases possibly lack signal peptide and are assumed to be localised in the cytoplasm. In general, as much as half of Fibrobacteres GHs are predicted to be localised in cytoplasm. At the same time, three times more GHs are exported outside the cell by Spirochaetae. This could indicate a rather selfish carbohydrates degradation strategy employed by Fibrobacteres. By maximizing intracellular cellulose breakdown they would avoid being in competition with other cellulolytic bacteria. Enrichment of *exbB* (encoding for a biopolymer transport protein ExbB; Fig. 2a) gene transcripts in Fibrobacteres metatranscriptome would suggest possible cellulose uptake through a system similar to a previously proposed TonB-dependent transport of maltodextrins across outer membrane of *Caulobacter crescentus*^35^.

## Discussion

Retrieving lignocellulose-active enzymes from naturally evolved biomass-degrading systems, with use of continuously improving high throughput sequencing technologies, presents a promising strategy to identify new enzymes with potentially enhanced activities. Limited metatranscriptomic reports highlight high representation and overexpression of CAZymes in termite digestomes (*e.g.* ref.^9,14^). In higher termite guts, many lignocellulolytic steps are assisted by gut microorganisms (*e.g.* cellulose deconstruction), with some being exclusively carried out by hindgut bacteria (*e.g.* hemicellulose degradation). Cellulose degradation capacities of different lignocellulose degrading environments, including the termite gut system^36^, have extensively been studied in the past. Decomposition of hemicellulose and general mobilisation of different lignocellulose components (breaking bonds between diverse plant polymers) have received comparably less scientific focus. Importantly, there is an increasing industrial interest in xylan-processing enzymes^37^, regarding their application in biomass (wood) processing, pulp bio-bleaching, animal nutrition, food additives, *etc.* (ref.^38^).

According to the recently established glycome profile of *Miscanthus* sp., next to glucose, xylose and arabinose are the two main cell wall monosaccharides, both originating from arabinoxylan fibres which are esther-cross-linked by ferulic acid ^39^. Consequently, *cazymes* specifically targeting (feruloylated) arabinoxylan components were highly up-regulated under *Miscanthus* diet, making them potentially suitable candidates for industrial xylan-targeting applications. Many of these genes were combined in clusters with a set of complementary hydrolytic activities to degrade *e.g.* AXs. Nature-optimised synergy between enzymes of the same CAZyme cluster could further provide the basis to better define industrially relevant enzymatic cocktails. Specific lignocellulose fractions could be selectively targeted to deliver desired products, with potential effects being fully expectable and controllable^40^. It would allow for their fine-tuned degradation for a variety of applications, *e.g.* oligoarabinoxylans for food industry (prebiotics), lignin fibres for biomaterials, glucomannans as food additives, *etc*. Feruloyl esterases, by removing cross-links between polysaccharides and lignin help separating lignin from the rest of biomass, offering an alternative and/or complementation to currently applied industrial treatments^41^. In addition, ferulic acid and other hydroxycinnamic acids can have many applications in food and cosmetic industries due to their antioxidant properties *etc.*^42^, thus further extending the application range of *Miscanthus* biomass.

Approach-wise, experimental design undertaken in this study represents the enrichment strategy where a nature-derived microbial inoculum is grown in liquid batch cultures. Here, a natural system of the termite gut was shown to progressively adapt yielding a consortium of microbes specialised in degradation of *Miscanthus* biomass. Integrative omics combined with protein characterisation provides a framework for better understanding of complex lignocellulose degradation by the higher termite gut system and paves a road towards its future bioprospecting.

## Methods

### Nest origin, laboratory maintenance and sampling

Initially, three colonies of grass-feeding higher termites from *Nasutitermitinae* (nests: LM1, LM2 and LM3) were identified in January 2017 in tropical savannah in French Guiana, in proximity to Sinnamary town (radius of 5 km to GPS: N 05°24.195’ W 053°07.664’). Termite nests were transported to the laboratory where colonies were maintained in separate glass containers at 26 °C, 12 h light and 12 h dark, and 90 % humidity conditions. Termite colonies were fed with *Miscanthus* grass winter harvest rich in recalcitrant lignocellulose^43^, for a period of up to nine months. Colony LM1_2 died after few months and was excluded from extended analysis. Mature worker-caste individuals were sampled in regular monthly time intervals before (sample taken before *Miscanthus* diet corresponds to “wild-microbiome”) and following *Miscanthus* diet (samples correspond to “*Miscanthus*-adapted microbiome”). Termite specimens were cold immobilized, surface-cleaned with 80 % ethanol and 1 x PBS and decapitated. Whole guts (here relative to the midgut and hindgut compartments) were dissected (n ≈ 30 per replicate, minimum three replicates per sample) and preserved directly in liquid nitrogen. Additionally, for a sample selected for metagenomics analysis (LM1 time point 8 months; LM1_8) the hindgut luminal fluid was collected as previously described^11^. Samples were stored at −80 °C until further processing. Termite species were identified by morphology and by sequencing of the partial COII marker gene, as described before^44^. The nucleotide sequences are available in GenBank under accession numbers MN803317-19.

### Extraction of nucleic acids

Total DNA and RNA were co-extracted from all samples using the AllPrep PowerViral DNA/RNA Kit (Qiagen) following the manufacturer’s protocol. To assure the proper disruption of bacterial cell wall and termite gut epithelium cells, mechanical bead-beating step with 0.1 mm glass beads at 20 Hz for 2 min was introduced to complement the chemical lysis. The eluents were divided in two parts. First part was treated with one µL of 10 µg/mL RNase A (Sigma) for 30 min at room temperature. The second part was treated with TURBO DNA-free kit (Invitrogen) according to manufacturer’s protocol. The resulting pure DNA and RNA fractions were quality assessed using agarose gel electrophoresis and Bioanalyser RNA 6000 Pico Kit (Agilent). Nucleic acid concentration was quantified using Qubit dsDNA HS Assay and Qubit RNA HS Assay Kit (Invitrogen). DNA and RNA were stored at −20 °C and - 80 °C, respectively.

### Bacterial 16S rRNA gene amplicon high-throughput sequencing and data analysis

The bacterial 16S rRNA gene amplicon libraries were prepared using Illumina compatible approach as previously described^45^. Briefly, modified universal primers S-D-Bact-0909-a-S-18 and S-*-Univ-*- 1392-a-A-15 (ref.^46^) and Nextera XT Index Kit V2 (Illumina) were used along with Q5 Hot Start High-Fidelity 2x Master Mix (New England Biolabs) to perform two-step polymerase-chain reaction (PCR). It allowed for selective amplification of the 484 bp long fragment of bacterial 16S rRNA gene V6–V8 region and simultaneous attachment of Illumina adapters and barcodes in the second step. Purified and equimolarly pooled libraries were sequenced along with PhiX control (Illumina) using MiSeq Reagent Kit V3-600 on in-house Illumina MiSeq Platform. Usearch v.7.0.1090_win64 software^47^ was used for quality trimming, chimera check, singletons removal and assignment of the obtained sequences to OTUs at 97 % similarity level. Taxonomic affiliation of the resulting OTUs was performed with SILVA database v.128 (ref.^48^). Raw sequences are available in the Sequence Read Archive (SRA) database under project number PRJNA587606. Resulting OTUs were deposited in GenBank under project numbers PRJNA586754 and PRJNA434195. Downstream analyses were performed with mothur^49^ and R environment^50^. Bacterial richness and diversity were calculated using respectively sobs and invsimpson indices. The dissimilarity of bacterial community structures was calculated using Bray-Curtis index. OTUs differentially abundant between the wild- and *Miscanthus*-adapted microbiomes were assessed using the LEfSe approach^12^.

### *De novo* metagenomics and data analysis

Sample LM1_8 was selected for metagenomic sequencing in order to reconstruct genomes/larger genomic fragments of the dominant microbes in the *Miscanthus*-adapted microbiome. Prokaryotic DNA was enriched from the total hindgut DNA extract using NEBNext Microbiome DNA Enrichment Kit (NewEngland BioLabs). Following sequencing, over 170 Mbp raw reads were quality trimmed in CLC Genomics Workbench v.9.5.2 (Qiagen), using a phred quality score of 20, minimum length of 50 and allowing no ambiguous nucleotides, resulting in close to 154 M quality-trimmed paired reads. Raw sequencing reads are available in the SRA database under project number PRJNA587423.

Quality-trimmed reads were assembled using the CLC’s *de novo* assembly algorithm in a mapping mode, using automatic bubble size and word size, minimum contig length of 1000, mismatch cost of 2, insertion cost of 3, deletion cost of 3, length fraction of 0.9, and similarity fraction of 0.95. The average contig abundance was calculated as DNA-RPKMs (reads per kilo base per million mapped reads). This type of normalization allows for comparing contigs (genomic fragments) coverage (abundance) values, as it corrects differences in both sample sequencing depth and contig length. ORFs were searched and annotated using the default pipeline integrated in the IMG/MER^51^.

Information about KEGGs and COGs assignment was retrieved based on the IMG/MER annotations. Metabolic pathways/modules were reconstructed using the tool integrated in the online version of the KEGG database^52^. Initially, contigs were binned using myCC^53^ what resulted in their separation into 35 population-level bins of relatively high contamination. This approach was undertaken to correctly assign phylum-level taxonomy to the resulting contigs. Subsequently, the bin refinement module integrated in MetaWRAP was used to fine-tune the resulting bins (MAGs) to the species/strain levels^54^. The completeness and contamination of the generated MAGs were assessed with checkM^55^. Taxonomic affiliation was assessed with PhyloPhlan^56^. Similarity to the previously reconstructed MAGs was verified with the FastANI^57^. MAGs abundance within the reconstructed metagenome was calculated as average of contigs metagenomic abundance (DNA-RPKMs, see above) assigned to a specific MAG. Given the relatively high microbial diversity in the termite gut, only 54.6 % of the resulting MG sequencing reads could map back to the reconstructed MG contigs, potentially mitigating the rate of functional gene discovery if solely relaying on the *de novo* MG reconstruction. Phylogenetic analyses were performed on MAFFT-aligned protein sequences^58^ using MEGA X^59^.

### *De novo* metatranscriptomics, host transcriptomics and data analysis

For three selected samples (LM1-1, LM1-2 and LM1-8) the (meta)transcriptomic analysis was performed using an optimised approach described earlier^11^. Ribo-Zero Gold rRNA Removal Kit “Epidemiology” (Illumina) was used to enrich the sample for prokaryotic and eukaryotic mRNA. Enriched mRNA was purified using Agencourt RNAClean XP Kit and analyzed with Bioanalyser RNA 6000 Pico Kit (Agilent). In continuation, SMARTer Stranded RNA-Seq Kit (Clontech) was used according to the manufacturer’s instructions to prepare sequencing libraries. Final libraries were quantified with High Sensitivity DNA Kit (Agilent) and KAPA SYBR FAST Universal qPCR Kit. Libraries were pair-end sequenced at the Luxembourg Centre for Systems Biomedicine (University of Luxembourg) using Illumina NextSeq 500/550 High Output v2-300 Kit. Raw sequencing reads are available in the SRA database under the project number PRJNA587406. Over 285 M raw reads were quality trimmed in CLC Genomics Workbench v.9.5.2, using a phred quality score of 20, minimum length of 50 and allowing no ambiguous nucleotides, resulting in close to 214 M quality-trimmed reads. Contaminating rRNA reads were removed using SortMeRNA 2.0 software^60^. The resulting non-rRNA reads were used to perform *de novo* (meta)transcriptome co-assembly using the CLC assembly algorithm in mapping mode with default parameters, except for minimum contig length of 200, length fraction of 0.90 and similarity fraction 0.95. As a result, nearly 2 M contigs were assembled. Obtained contigs were further submitted to IMG/MER for taxonomic and functional annotation^51^. Following the taxonomic assignment, 759,451 transcripts of putative prokaryotic origin were selected for further analysis. Initial IMG/MER taxonomic annotation resulted in over-representation of transcripts of putative Firmicutes origin (Supplementary Fig. 4). As nearly no Firmicutes OTUs were detected using the 16S rRNA gene amplicon sequencing, transcripts were re-annotated based on the *de-novo* assembled metagenome and contig binnig, resulting in re-classification of virtually all Firmicutes-assigned contigs to Fibrobacteres and Spirochaetae. To complement the study and to characterise potential contribution of the termite host to *Miscanthus* digestion, transcripts of eukaryotic origin and taxonomically assigned to *Insecta* (based on the IMG/MER annotation) were further evaluated for the completeness of the *de novo* reconstructed transcriptome with the BUSCO pipeline^26^ and using the Eukaryota database (odb9).

For both the *de novo* assembled metatranscriptome and termite transcriptome, in order to determine the relative abundances of transcripts across studied samples, sequencing reads were mapped back to the annotated transcript sets, using the CLC “RNA-seq analysis” mode, with default parameters except for minimum similarity of 0.95 over 0.9 of the read length, both strands specificity and 1 maximum number of hits per read. The mapping results were represented as TPMs (transcripts per million), what directly resulted in normalised reads counts.

### Identification of *cazymes*, heterologous protein production, purification and activity testing

Genes (metagenomics) and gene transcripts (metatranscriptomics) encoding for carbohydrate active enzymes were detected using dbCAN (dbCAN-fam-HMMs.txt.v6; ref.^61^) and CAZy database^62^. Using the threshold of e-value of < 1e^-18^ and coverage > 0.35 recommended for prokaryotic CAZymes resulted in removal of a high number of putative CAZymes, therefore additional manual curation was performed to maximise the number of entries retained for further analysis. Additionally, gene transcripts outliers (very partially reconstructed gene fragments with average MT expression significantly exceeding the average expression of other genes assigned to the same group) were manually identified and removed as they were considered as chimeric (additional Blast search was launch in each case). Homology to peptide pattern (Hotpep) was used to assign an EC class to the identified CAZymes (ref. ^21^). Sub-cellular localisation of CAZymes was predicted using BUSCA web^63^. To link the degradation of the different lignocellulose fractions and subsequent sugar utilisation, we looked for the presence of suitable sugar transporters and also specific sugar isomerases and kinases.

Genes encoding for CAZymes of interest, selected based on their predicted activities and their expression profiles, were further PCR amplified (Veriti™ 96 wells Thermal cycler, Applied Biosystems, Foster City, USA) and the resulting PCR products were purified using a PCR purification kit (Qiagen, Hilden, Germany). If any signal peptide was predicted (using LipoP version 1.0, http://www.cbs.dtu.dk/services/LipoP/, ref.^64^), it was removed before cloning to enhance cytoplasmic protein production. Purified PCR products were cloned into the pET52b(+) plasmid and expressed in *E. coli* Rosetta (DE3) strain (Millipore Corporation, Billerica, MA, USA), as previously described^40^. Cells were harvested by centrifugation (5,000 x g, 4°C, 15 min) and re-dissolved in a lysis buffer (50 mM NaH_2_PO_4_, 10 mM imidazole, 300 mM NaCl, pH8). Proteins were released by sonication, cell debris were removed by centrifugation (16,000 x g, 4°C, 15 min) and subsequent filtration step (13-mm syringe filter, 0.2-µm-pore-size). Affinity tag purification was achieved using a histidine tag located at the C terminus of a recombinant protein. NGC™ Medium-Pressure Liquid Chromatography system (Bio-Rad) equipped with a Hitrap™ column of 5 mL (Bio-Rad, Hercules, CA, USA) was used to purify produced proteins. A constant flow rate of 5mL/min was applied. Initially, column was equilibrated with six column volumes (CV) of lysis buffer. Following equilibration, 290 mL of sample was injected and washed with three CVs of mixed buffer (97 % of lysis buffer and 3 % of elution buffer, the later composed of 50 mM NaH_2_PO_4_, 250 mM imidazole, 300 mM NaCl, pH8).

First step of elution was achieved using ten CVs of a linear gradient of mixed buffer (from 3 % of elution buffer plus 97 % of lysis buffer to 50 % of elution buffer plus 50 % of lysis buffer), followed by four CVs of 100 % of elution buffer and finally one CV of 100 % lysis buffer to detach the remaining protein. Fractions of two mL were collected during the washing and elution steps, and they were analysed on SDS-PAGE. The release of 4-nitrophenol (PNP assay) and/or reducing sugar (RS assay) was used to determine the activity of recombinant proteins. Briefly, 50 µL of a purified protein solution was incubated with 50 µL of substrate (respectively, 4-nitrophenol derivatives were used for PNP assay and polysaccharides for RS assay) and 100 µL (PNP assay) or 25 µL (RS assay) of citrate phosphate buffer pH7 (0.1M citric acid, 0.2M dibasic sodium phosphate). The targeted substrates included carboxymethylcellulose (CMC), arabinoxylan, galactomannan, glucomannan and xylan. Enzymatic reaction was carried out at 37 °C during one hour (PNP assay) or 30 min (RS assay). The rate of release of 4-nitrophenol was instantly monitored at 405 nm. The release of reducing sugars was determined following the Somogyi-Nelson method (ref.^65,66^). All assays were performed in triplicates.

## Acknowledgements

This research was funded through FNR 2014 CORE project OPTILYS (Exploring the higher termite lignocellulolytic system to optimize the conversion of biomass into energy and useful platform molecules/C14/SR/8286517), and co-funded through the grant PDR T.0065.15 from the Belgian F.R.S.-FNRS. We are grateful to Philippe Cerdan, Régis Vigouroux and the staff of the Laboratoire Environnement HYDRECO of Petit Saut (EDF-CNEH) for logistic support during the field work.

## Author contributions

Experiments and molecular analyses were planned by MC, MM, MB and DSD and carried out by MC, MM and MB with a technical support from BU and DK. MC and MM analysed the results. MB performed protein studies. YR, DSD, XG helped with the collection of termites. RH and PW provided the support with Illumina NextSeq sequencing. PG and XG provided support with bioinformatics analysis. MC, MM and MB wrote the manuscript. PD, YR, DSD, PG, PF participated in the planning and coordination of the study and in the manuscript correction. All authors read and approved the final manuscript.

## Additional information

Supplementary information is provided for this paper as separate file and tables.

