## Supplementary_material for "Targeted biomass degradation by the higher termite gut system - integrative omics applied to host and its gut microbiome"

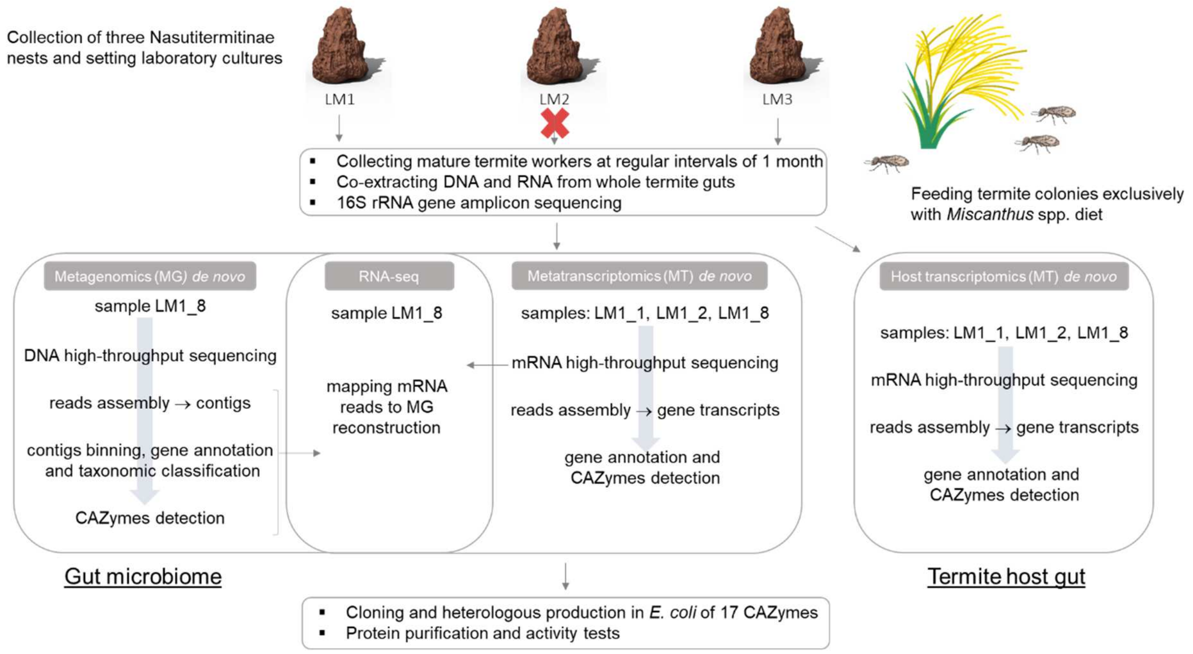


**Supplementary Figure 1**: Assay design of sampling and high-throughput sequencing characterisation of the termite gut lignocellulose digestion system. Termite hindguts from mature workers were sampled in regular monthly time intervals and nucleic acids were co-extracted. Colony LM2 was excluded from further analysis as it did not adapt to the laboratory fed *Miscanthus* spp. and diet. Control sample (fed original diet) is designated as LMx_1. Both, the termite gut microbiome (bacterial community) and the termite gut epithelium were analysed.

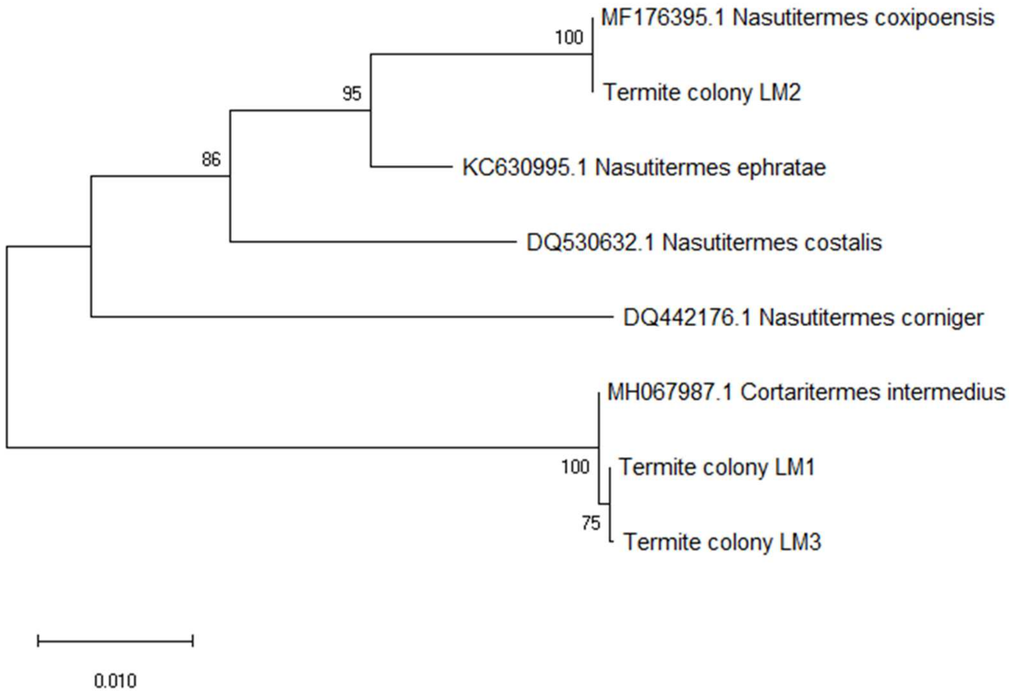


**Supplementary Figure 2:** Unrooted neighbor-joining tree of cytochrome oxidase II genes of the three termite species investigated in this study (LM1, LM2 and LM3) and their closest sequenced relatives, based on the homology search against the NCBI database. The percentage of replicate trees in which the associated taxa clustered together in the bootstrap test (500 replicates) are shown next to the branches. There were a total of 747 positions in the final dataset. Evolutionary analysis was conducted in MEGA X[1].

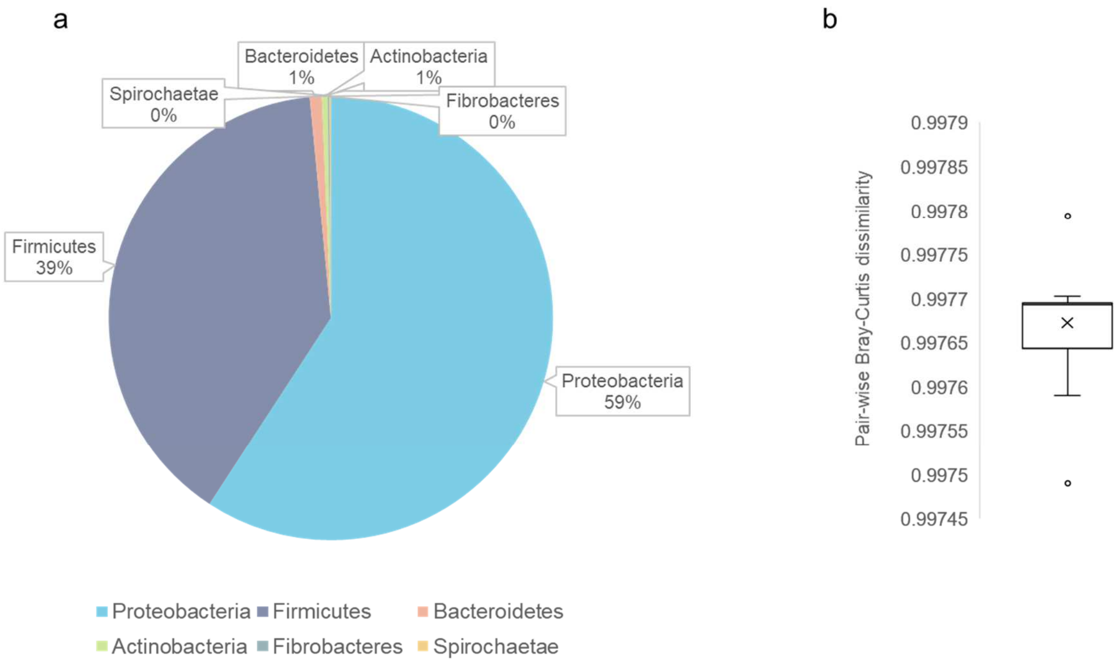


**Supplementary Figure 3**: Characterisation of the *Miscanthus* straw associated bacteria. (a) Taxonomic distribution of the 16S rRNA reads to bacterial phyla. (b) Pair-wise Bray-Curtis dissimilarity between the *Miscanthus* straw associated microbial community and the *Miscanthus*-adapted microbiome (bacterial community in the termite gut fed with *Miscanthus* diet). The lowest calculated pair-wise Bray-Curtis dissimilarity distance was above 0.997 (on the scale from 0 to 1, where 0 means that two communities are identical and 1 means that they are maximally different), and none of the *Miscanthus* straw associated microbes was enriched in the termite gut microbiome.

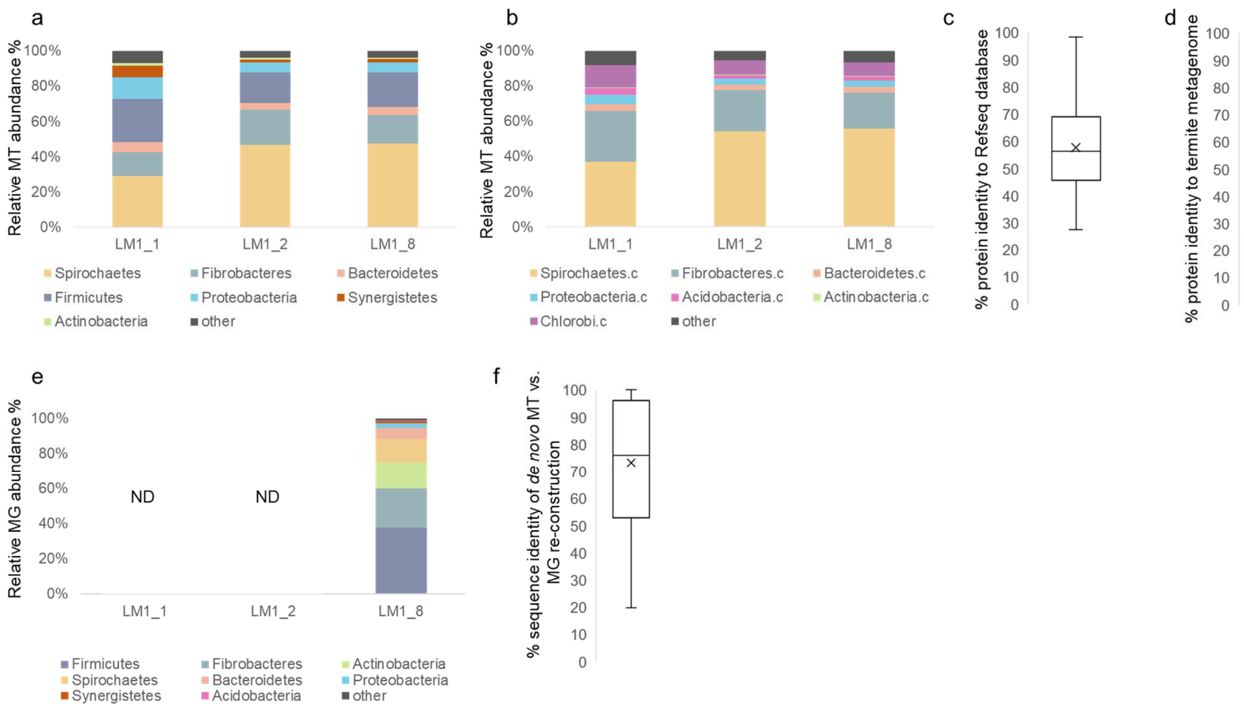


**Supplementary Figure 4**: (a) Database-dependent (IMG/MER; ref.[2]) taxonomic assignment of reconstructed *de novo* MT gene transcripts for the studied termite gut microbiomes. (b) Taxonomic reclassification of the *de novo* MT reconstructed gene transcripts for the studied termite gut microbiomes based on their sequence homology to the *de novo* MG reconstructed contigs and bacterial MAGs. (c, d) Sequence similarity comparison of the *de novo* reconstructed gene transcripts for the studied termite gut microbiomes against the NCBI nt database (c) and a custom *Nasutitermes* spp database (d; ref.[3]). (e) Database-dependent (IMG/M; ref.[2]) taxonomic assignment of reconstructed *de novo* MG genes for the studied termite gut microbiome. (f) Comparison of the de novo MG and MT reconstructions, based on the sequence similarity of the reconstructed genes (MG) versus gene transcripts (MT). ND – sequencing not done.

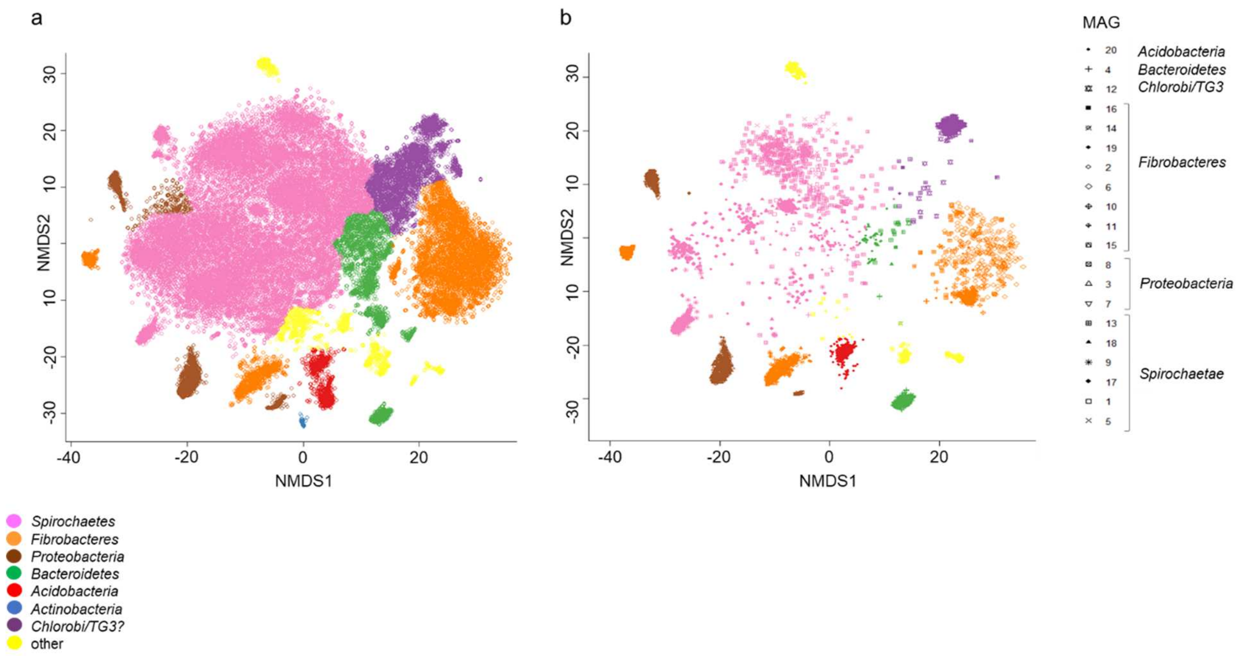


**Supplementary Figure 5:** (a) Binning results of the *de novo* reconstructed metagenomic contigs to phylum-level bins and taxonomic bin assignment with PhyloPhlan[4]. (b) Reconstruction of species-level MAGs (metagenome assembled genomes) and their phylogenetic classification.

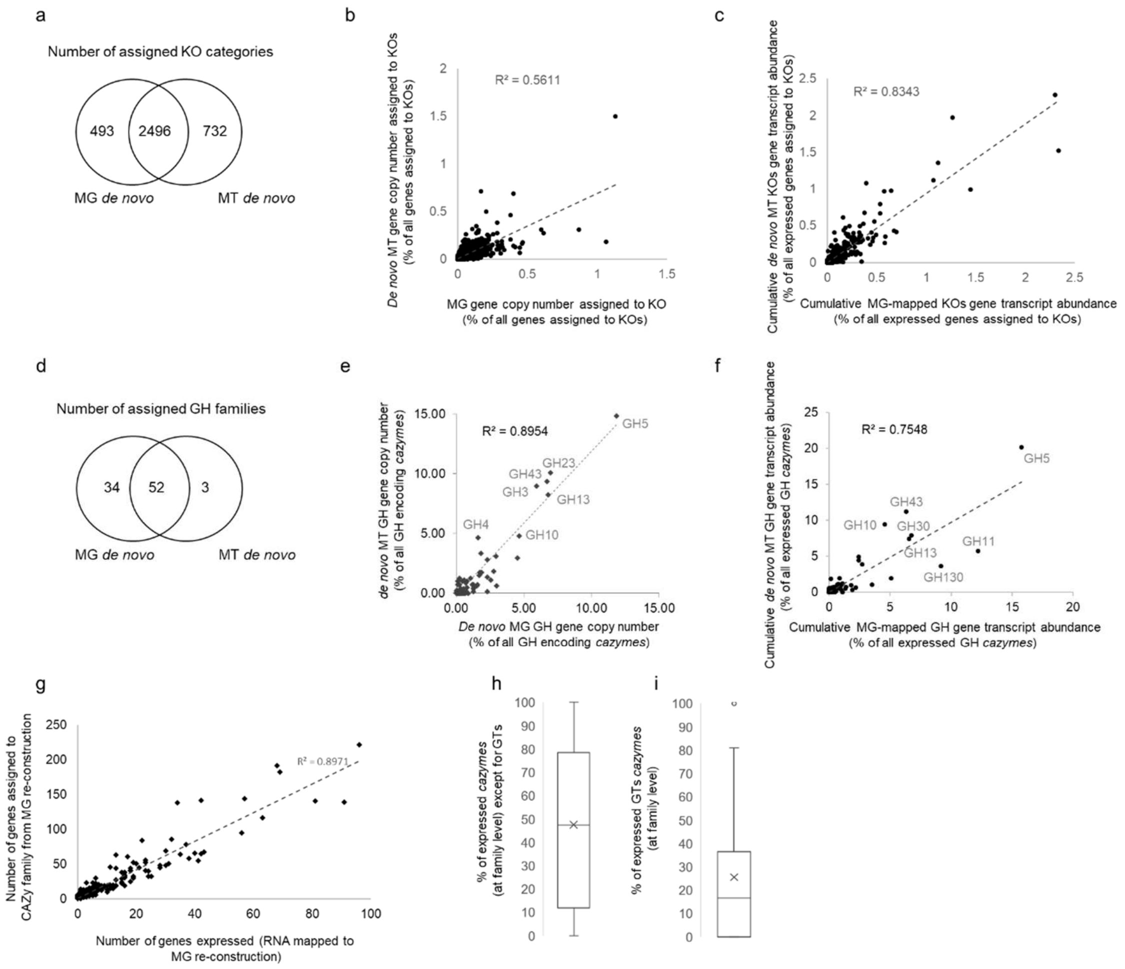


**Supplementary Figure 6:** Comparison of the *de novo* MG and MT reconstructions for the studied termite gut microbiomes. (a) Venn diagram showing unique and shared KOs between the two datasets. (b) Comparison of the number of reconstructed genes (MG) and mapped gene transcripts (MT) assigned to the same KO category for sample LM1_8. Pearson coefficient of correlation is displayed on the graph. (c) Comparison of the cumulative gene (MG) and gene transcript (MT) abundance assigned to the same KO category. Pearson coefficient of correlation is displayed on the graph. (d) Venn diagram showing unique and shared GH families between the two datasets. (e) Comparison of the number of reconstructed genes (MG) and gene transcripts (MT) assigned to the same GH family. Pearson coefficient of correlation is displayed on the graph. (f) Comparison of the cumulative gene (MG) and mapped gene transcripts (MT) abundance assigned to the same GH family for sample LN1_8. Pearson coefficient of correlation is displayed on the graph. (g) Comparison of the number of assigned genes (MG) to GH families and expressed genes (MT) assigned to the same GH family for sample LN1_8. Pearson coefficient of correlation is displayed on the graph. (h, i) Box plots representation with the median, first and third quartiles displayed of the average gene number expression per CAZy family, with a separate focus on glycosyl transferases GTs (i).

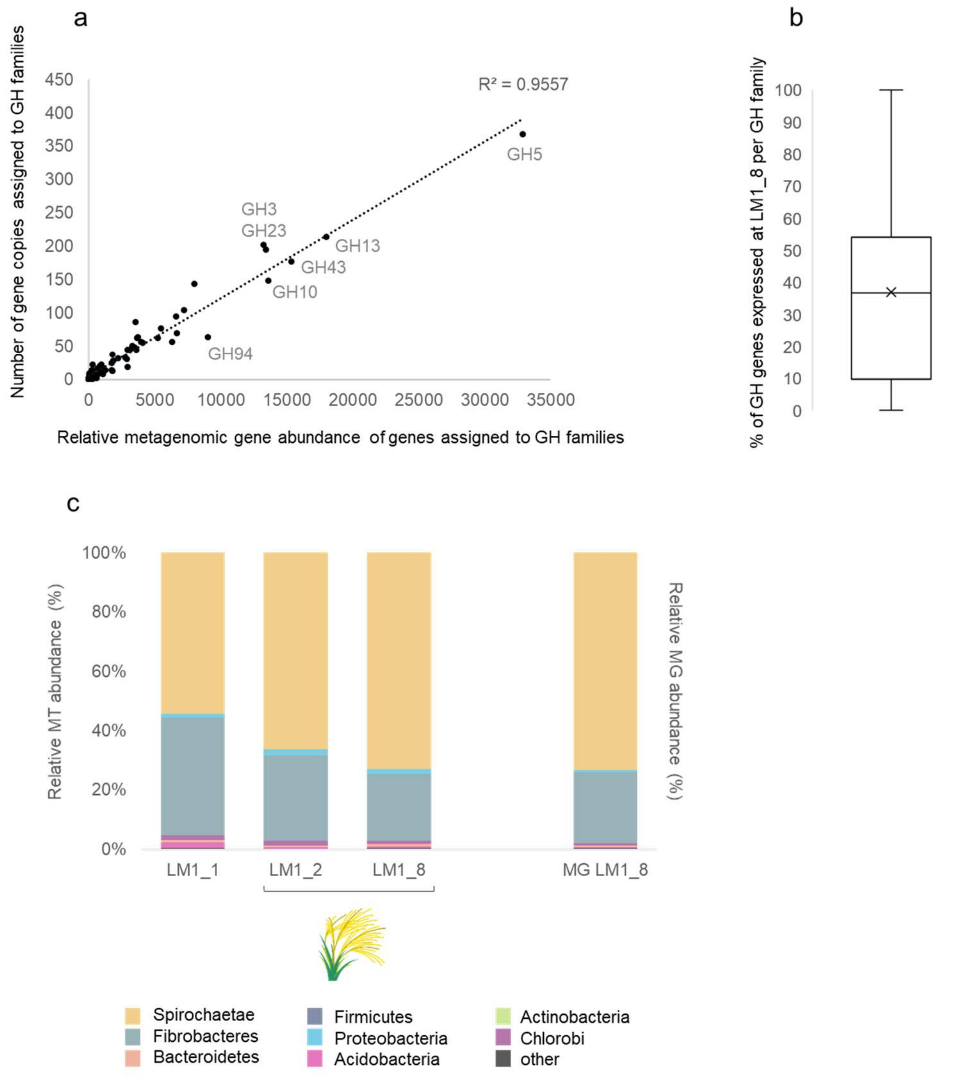


**Supplementary Figure 7:** Characterisation of microbial genes assigned to the different GHs families for the *de novo* MG reconstruction. (a) Correlation between the number of genes assigned to a GH family and their cumulative MG abundance. In the case of the GH11 family, the highly abundant and partially reconstructed gene outliers were not displayed on the graph. Pearson coefficient of correlation is displayed on the graph. (b) Number of the *de novo* MG reconstructed genes assigned to the different GH families that were expressed at the time point LM1_8; given per family (RNA-seq). (c) Database-independent classification of the *de novo* reconstructed genes (MG) and gene transcripts (MT) to the phylum level, and based on the MG contig binning and bin taxonomic annotation with PhyloPhlan[4].

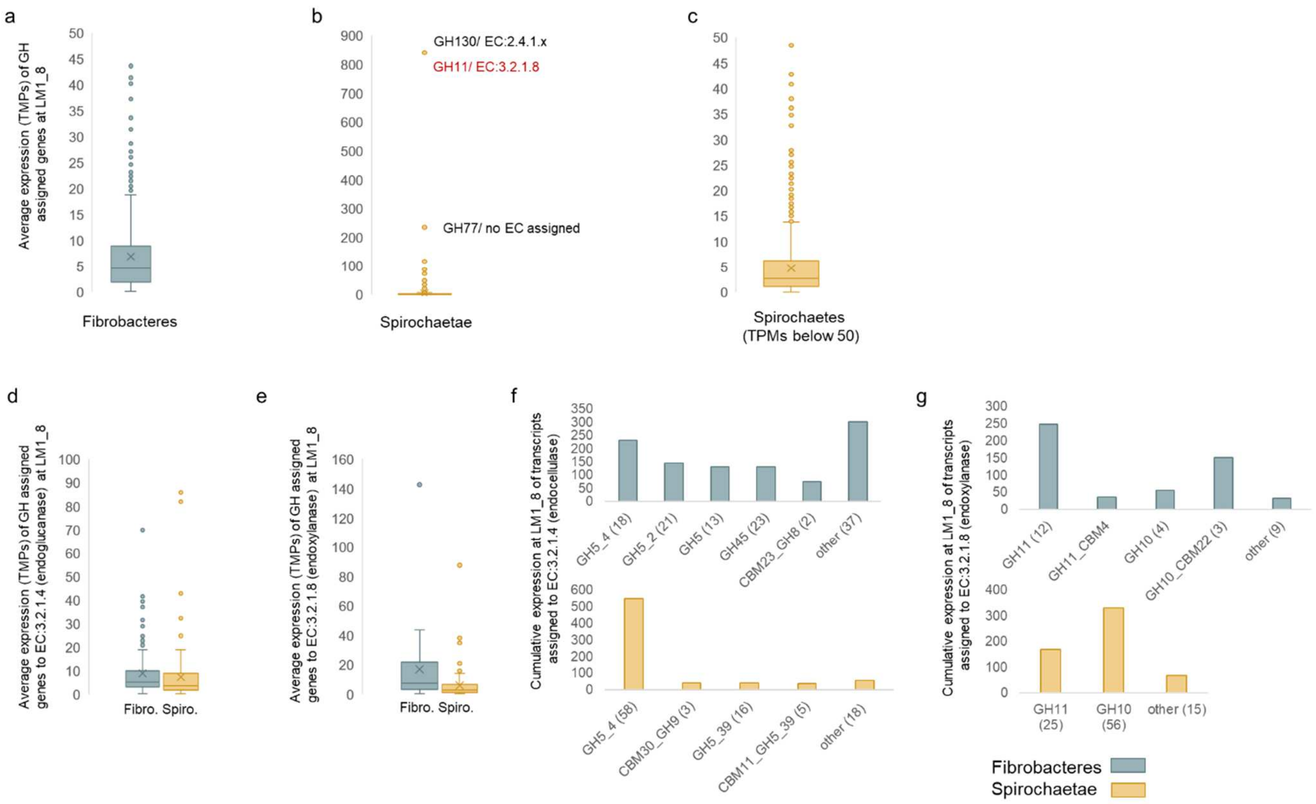


**Supplementary Figure 8:** Phylum level (Spirochaetae and Fibrobacteres) characterisation of the termite gut microbial GH coding genes and their expression profiles. Average MT abundance of gene transcripts assigned to the different GH families for (a) Fibrobacteres and Spirochaetae (b, c). Average MT abundance of GH gene transcripts functionally assigned to endoglucanases (d) and endoxylanases (e). Cumulative MT abundance of GH gene transcripts functionally assigned to endoglucanases (f) and endoxylanases (g). Distribution of gene transcripts between the different GH families is shown as bar charts. Number of reconstructed gene transcripts is shown in brackets. Gene transcripts outliers (highly abundant but partially reconstructed gene transcripts) were removed from panels c, e and g.

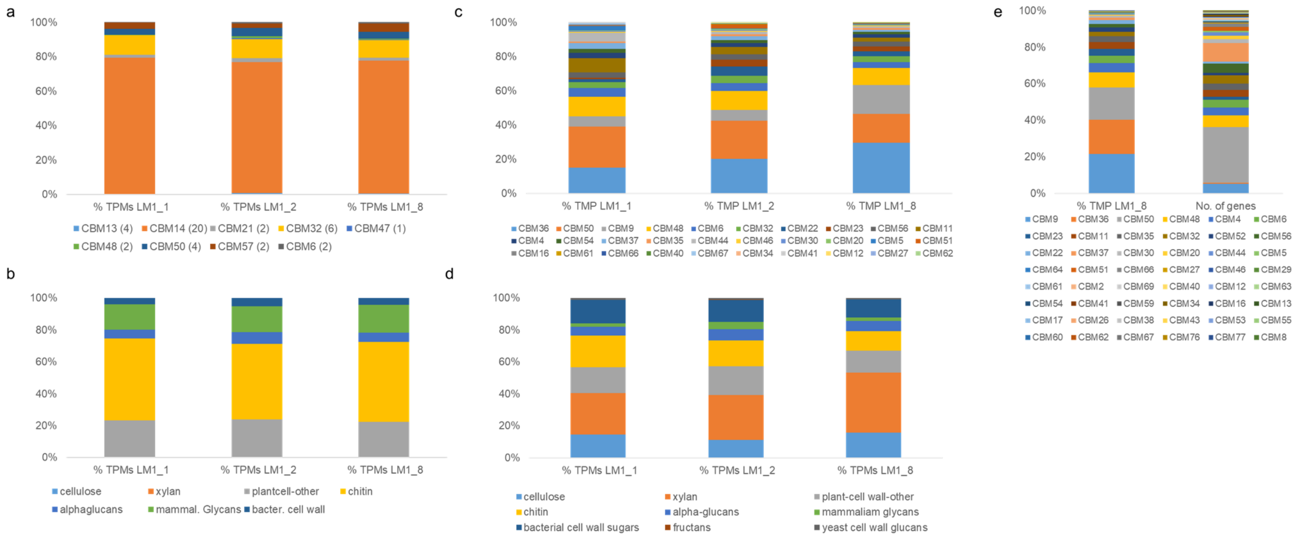


**Supplementary Figure 9:** Characterisation of termite and microbial genes (*de novo* MG) and gene transcripts (*de novo* MT) assigned to the different CBM families. (a) Distribution of reconstructed gene transcripts of termite origin to CBM families. Total number of genes is given in brackets. (b) Substrate specificity of reconstructed CBM encoding genes of termite origin, based on known substrate specificities of different CBM families. (c) Distribution of reconstructed gene transcripts (*de novo* MT) of microbial origin to CBM families. (d) Substrate specificity of reconstructed CBM encoding genes of microbial origin, based on known substrate specificities of different CBM families (e) Metagenomic abundance and gene number of the different CBM families, based on the annotation of the *de novo* reconstructed genes (*de novo* MG). For all graphs, the data is expressed as % of total CBM gene transcript abundance or gene count. Note that the colour code may change between the different panel in this figure.

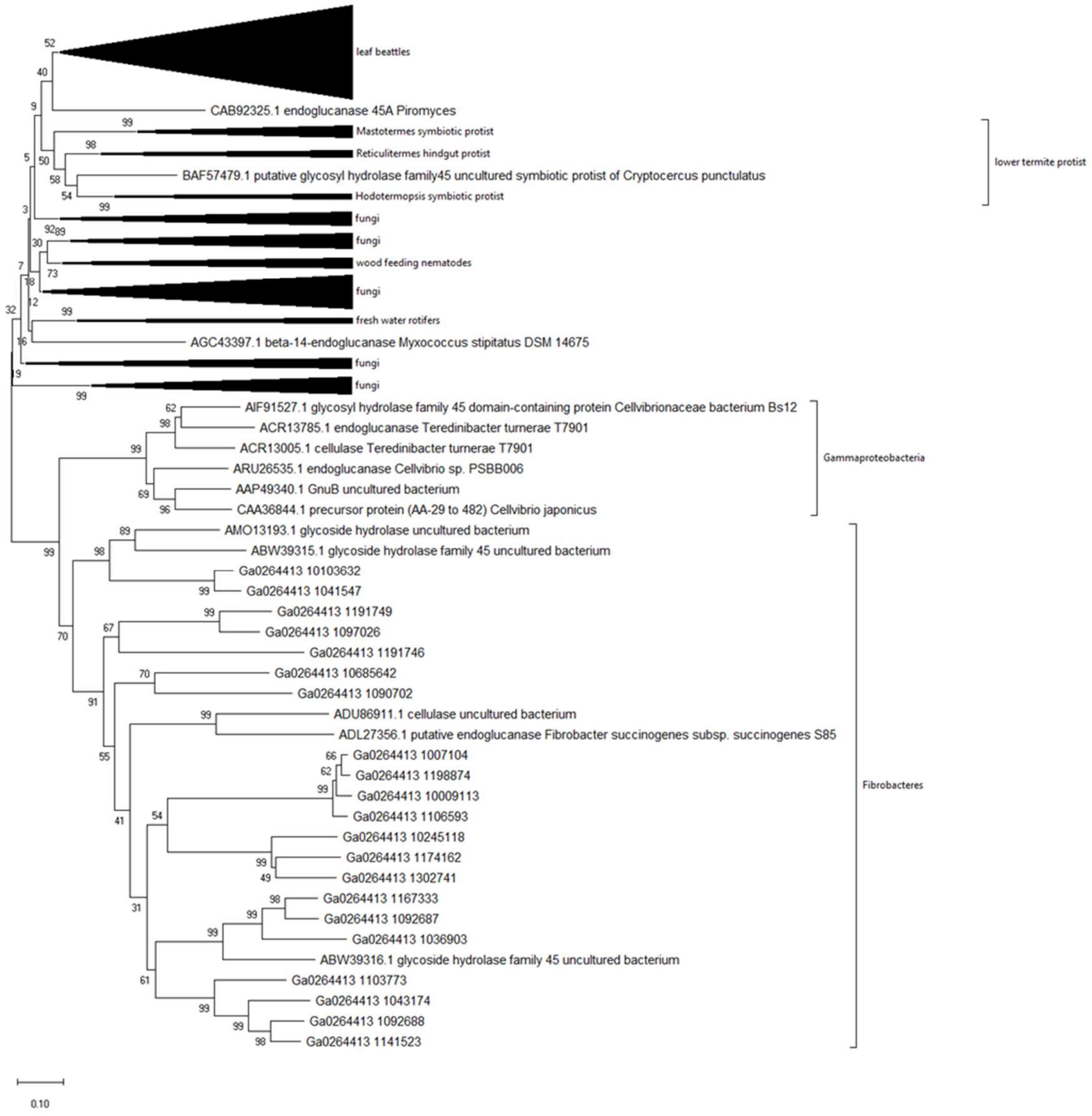


**Supplementary Figure 10:** Unrooted neighbor-joining tree of GH45 assigned genes, based on the *de novo* MG reconstruction. All known bacterial and eukaryotic genes assigned to GH45 in the CAZY database (http://www.cazy.org) are included. Some short sequences were removed from final alignment to increase the number of final positions in the alignment that could have been compared. The percentage of replicate trees in which the associated taxa clustered together in the bootstrap test (500 replicates) are shown next to the branches. There were a total of 486 positions in the final dataset. Evolutionary analysis was conducted in MEGA X[1].

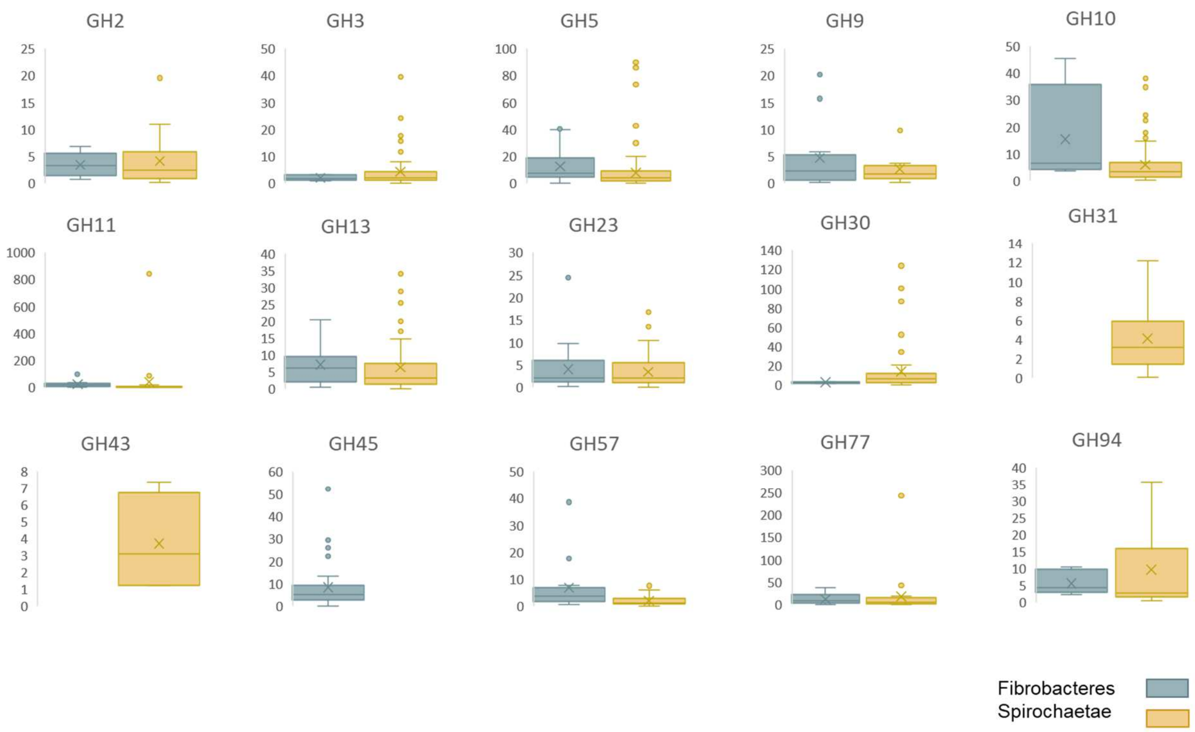


**Supplementary Figure 11:** Average per gene expression (TMP) for Spirochaetae and Fibrobacteres assigned GH genes (RNA-seq results) at LM1_8. Box plots representation with the median, first and third quartiles is displayed. Outliers (highly expressed genes; in some case representing only partially reconstructed genes) are indicated on the different panels with a dot.

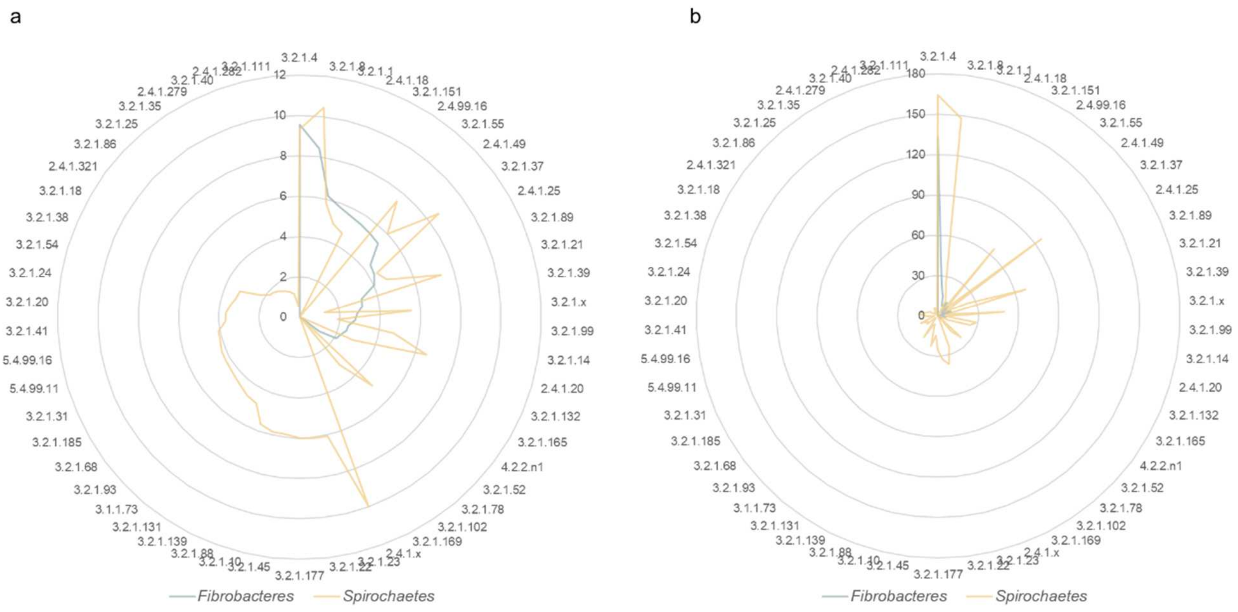


**Supplementary Figure 12**: Distribution of all GH-assigned genes of Fibrobacteres and Spirochaetae origin to different EC categories. (a) Relative transcriptional abundance (expressed as TPMs, log2 transformed) of reconstructed gene transcripts (*de novo* MT) at LM1_8. (b) Number of gene copies.

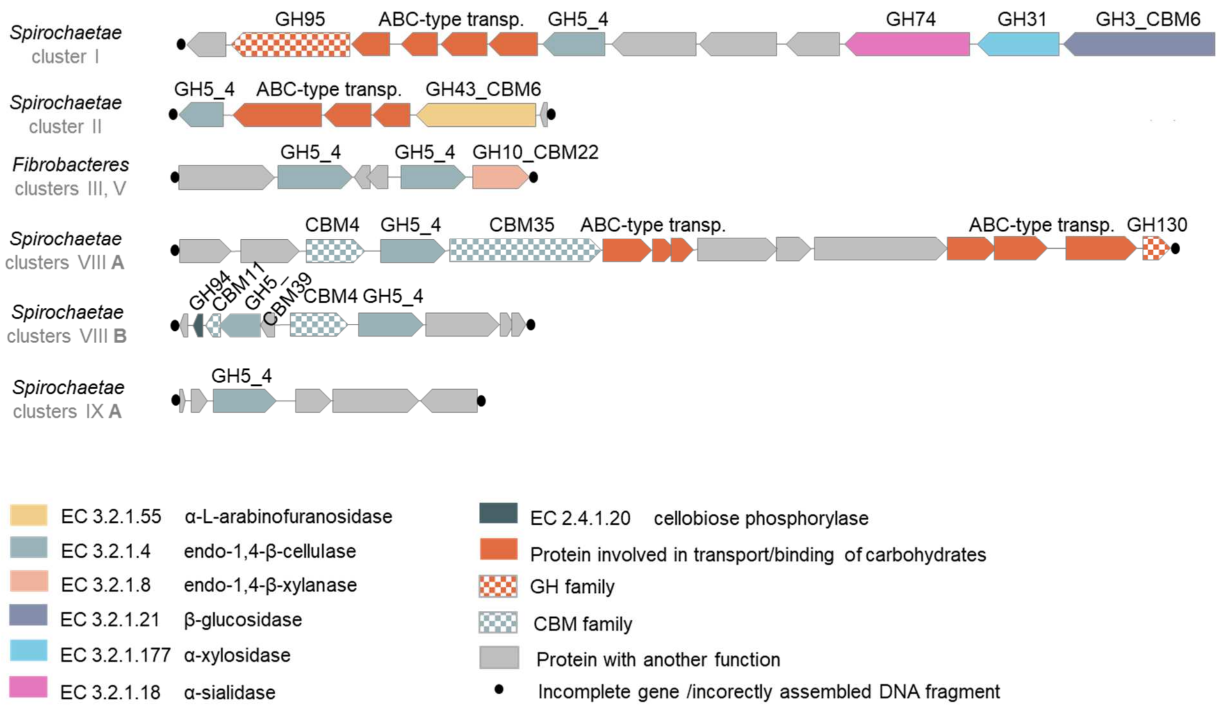


**Supplementary Figure 13:** Schematic representation of gene organisation within putative CAZymes genes loci. Genomic reconstructions are partial, therefore the CAZymes clusters may be fragmented (incomplete). Clusters designation (I to IX) refers to the Fig. 5a in the main text.

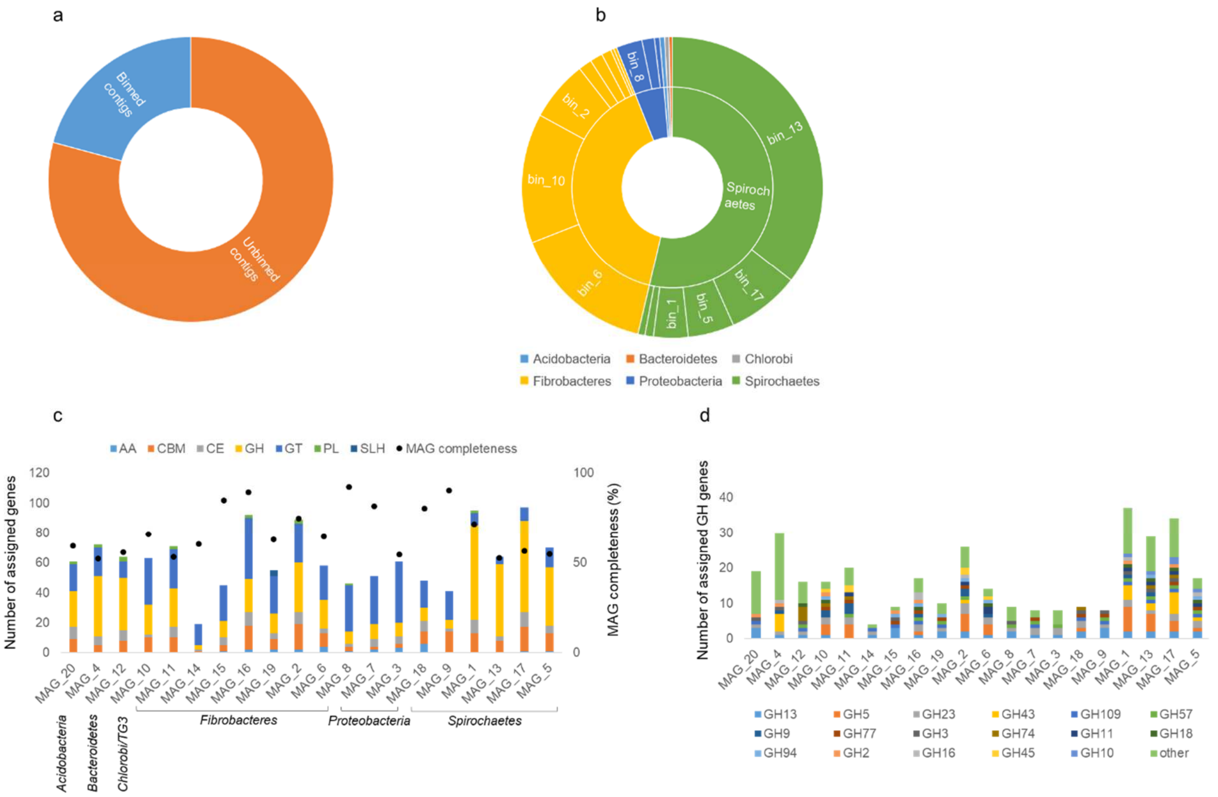


**Supplementary Figure 14:** Characterisation of reconstructed MAGs. (a) Proportion of reconstructed contigs that were further binned to MAGs, out of the whole *de novo* MG reconstruction for sample LM1_8. (b) Metagenomic abundance of reconstructed MAGs (proportion of all MAGs) based on the average abundance of reconstructed contigs binned into specific MAGs. MAGs are coloured according to their phylum-level taxonomic assignment. (c) CAZymes content of the reconstructed MAGs. (d) Diversity of the dominant GHs present in the reconstructed MAGs.

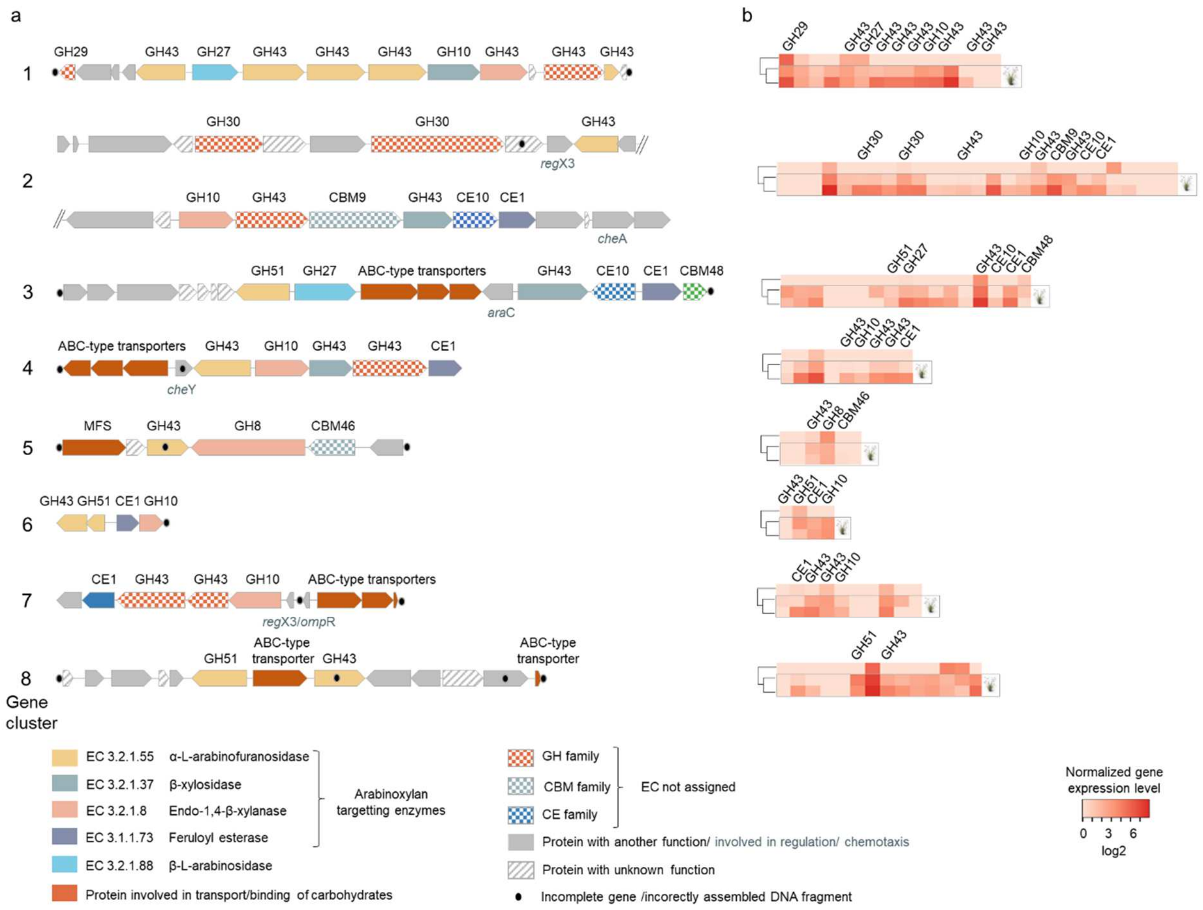


**Supplementary Figure 15:** Schematic representation of gene organisation within putative arabinoxylan-targeting CAZymes clusters of Spirochaetae origin (a). Genomic reconstructions are partial, therefore the CAZymes clusters may be fragmented. (b) Gene expression levels before and under *Miscanthus* diet for genes indicated in panel a.

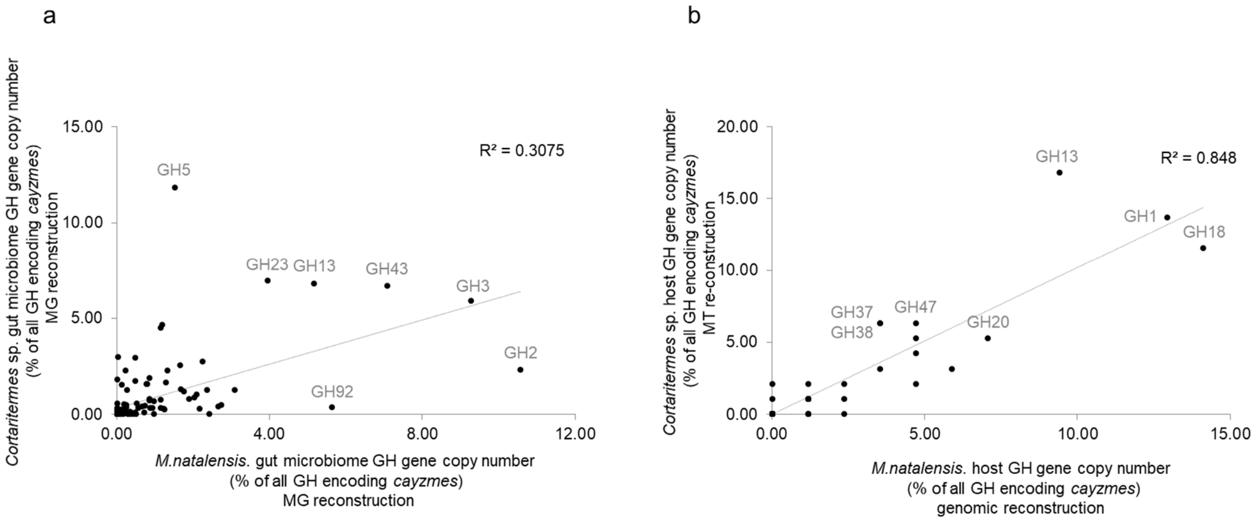


**Supplementary Figure 16:** Comparison of the GH content of the studied *Cortaritermes* gut system with a previously studied *M. natalensis*[5]. (a) Comparison of the GH gene diversity between the gut microbiomes of the two studied species. (b) Comparison of the GH gene diversity between the two studied termite species.

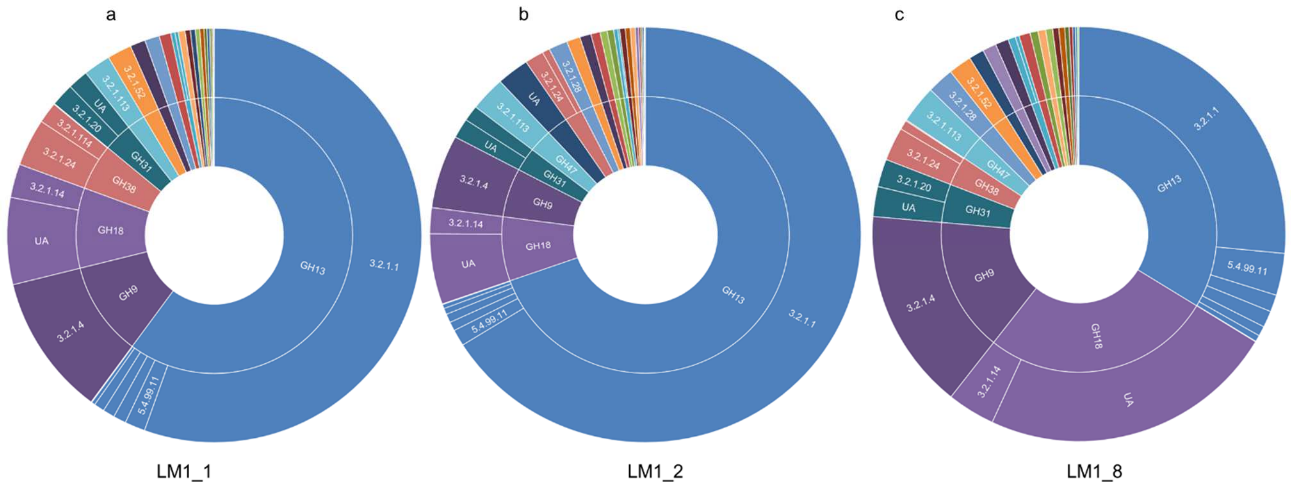


**Supplementary Figure 17:** Distribution of GH-assigned gene transcripts of *Cortaritermes* sp. origin to different EC categories. (a) Relative transcriptional abundance (expressed as TPMs, log2 transformed) of reconstructed gene transcripts (*de novo* MT) at LM1_1; (b) at LM1_2; and (c) at LM1_8.

**Supplementary Guide**

**Supplementary Table 1:** Details of the 16S rRNA gene amplicon high-throughput sequencing. Data is normalised to 10,000 reads per sample. Technical replicates are indicated with the same sample name.

**Supplementary Table 2:** Linear discriminant analysis (LDA) effect size (LEfSe; Segata *et al*., 2011) of bacterial OTUs significantly enriched in wild-microbiome (bacterial population in a termite gut feeding natural food) and in *Miscanthus*-adapted microbiome (bacterial population in a termite gut feeding on *Miscanthus* spp. grass)

**Supplementary Table 3:** Relative abundance of genes (metagenomics) and gene transcripts (metatranscriptomics, host transcriptomics) assigned to Clusters of Orthologous Groups (COGs) for gut microbiome and host transcriptome.

**Supplementary Table 4:** KEGG ontology profiles (KOs) assigned to genes (*de novo* MG) and gene transcripts (*de novo* MT) of prokaryotic (gut microbiome) and eukaryotic (termite host) origin.

**Supplementary Table 5:** Linear discriminant analysis (LDA) effect size (LEfSe; Segata *et al*., 2011) of KOs significantly enriched in the two main bacterial populations, Fibrobacteres and Spirochaetae, based on the RNA-seq results. Metabolic modules (only complete and with one block missing) reconstruction for the two main bacterial populations in the termite gut, Spirochaetae and Fibrobacteres, based on the RNA-seq results.

**Supplementary Table 6:** Characterisation of heterologously produced CAZymes.

**Supplementary Table 7:** Characteristics of the reconstructed metagenome assembled genomes (MAGs) based on the checkM evaluation. Putative CAZymes and CAZymes genes clusters on the *de novo* reconstructed MG contigs.

**Supplementary Table 8:** Transporters reconstruction, comparison of the *de novo* metagenomics, *de novo* metatranscriptomics and RNA-seq for the termite gut microbiome.

**Supplementary Table 9:** Comparison of CAZy genomic (cazymes domains; here defined based on the expressed termite host *cazymes*) content between the different termite species with sequenced genomes).

**Supplementary Table 10:** Per phylum summary of the sub-cellular localisation of GH-assigned genes and further functionally annotated with HotPep.
